# Next generation APOBEC3 inhibitors: Optimally designed for potency and nuclease stability

**DOI:** 10.1101/2024.09.05.611238

**Authors:** Adam K. Hedger, Wazo Myint, Jeong Min Lee, Diego Suchenski-Loustaunau, Vanivilasini Balachandran, Ala M. Shaqra, Nese Kurt-Yilmaz, Jonathan K. Watts, Hiroshi Matsuo, Celia A. Schiffer

## Abstract

APOBEC3 (or A3) enzymes have emerged as potential therapeutic targets due to their role in introducing heterogeneity in viruses and cancer, often leading to drug resistance. Inhibiting these enzymes has remained elusive as initial phosphodiester (PO) linked DNA based inhibitors lack stability and potency. We have enhanced both potency and nuclease stability, of 2′-deoxy-zebularine (dZ), substrate-based oligonucleotide inhibitors for two critical A3’s: A3A and A3G. While replacing the phosphate backbone with phosphorothioate (PS) linkages increased nuclease stability, fully PS-modified inhibitors lost potency (1.4-3.7 fold) due to the structural constraints of the active site. For both enzymes, mixed PO/PS backbones enhanced potency (2.3-9.2 fold), while also vastly improving nuclease resistance. We also strategically introduced 2′-fluoro sugar modifications, creating the first nanomolar inhibitor of A3G-CTD2. With hairpin-structured inhibitors containing optimized PS patterns and LNA sugar modifications, we characterize the first single-digit nanomolar inhibitor targeting A3A. These extremely potent A3A inhibitors, were highly resistant to nuclease degradation in serum stability assays. Overall, our optimally designed A3 oligonucleotide inhibitors show improved potency and stability, compared to previous attempts to inhibit these critical enzymes, opening the door to realize the therapeutic potential of A3 inhibition.

## INTRODUCTION

APOBEC3 (or A3) are a series of seven human and primate enzymes that play a key role both in the innate immune response to invading viruses (1,2) and in a variety of cancers (3–5). These enzymes catalyze the deamination of cytidine (C) to uridine (U) in single-stranded DNA (ssDNA) substrates (**Fig 1A**). In HIV-1 infection, where they were first characterized, A3G mutates the viral genome to such an extent that the virus has evolved to express a factor called Vif that targets certain A3 enzymes including APOBEC3D, F, G and H for proteasomal degradation (6). Nevertheless, the mutational signatures of APOBEC3G (A3G) have been associated with viral drug resistance and immune escape (7,8). In a broad range of cancers, genome variation has been linked to certain A3 enzymes, particularly A3A (3,8–13) (14), which leads to drug resistance to many therapies. Therefore, it is of significant interest to develop inhibitors of A3 enzymes as potential therapeutics.

**Figure 1.**
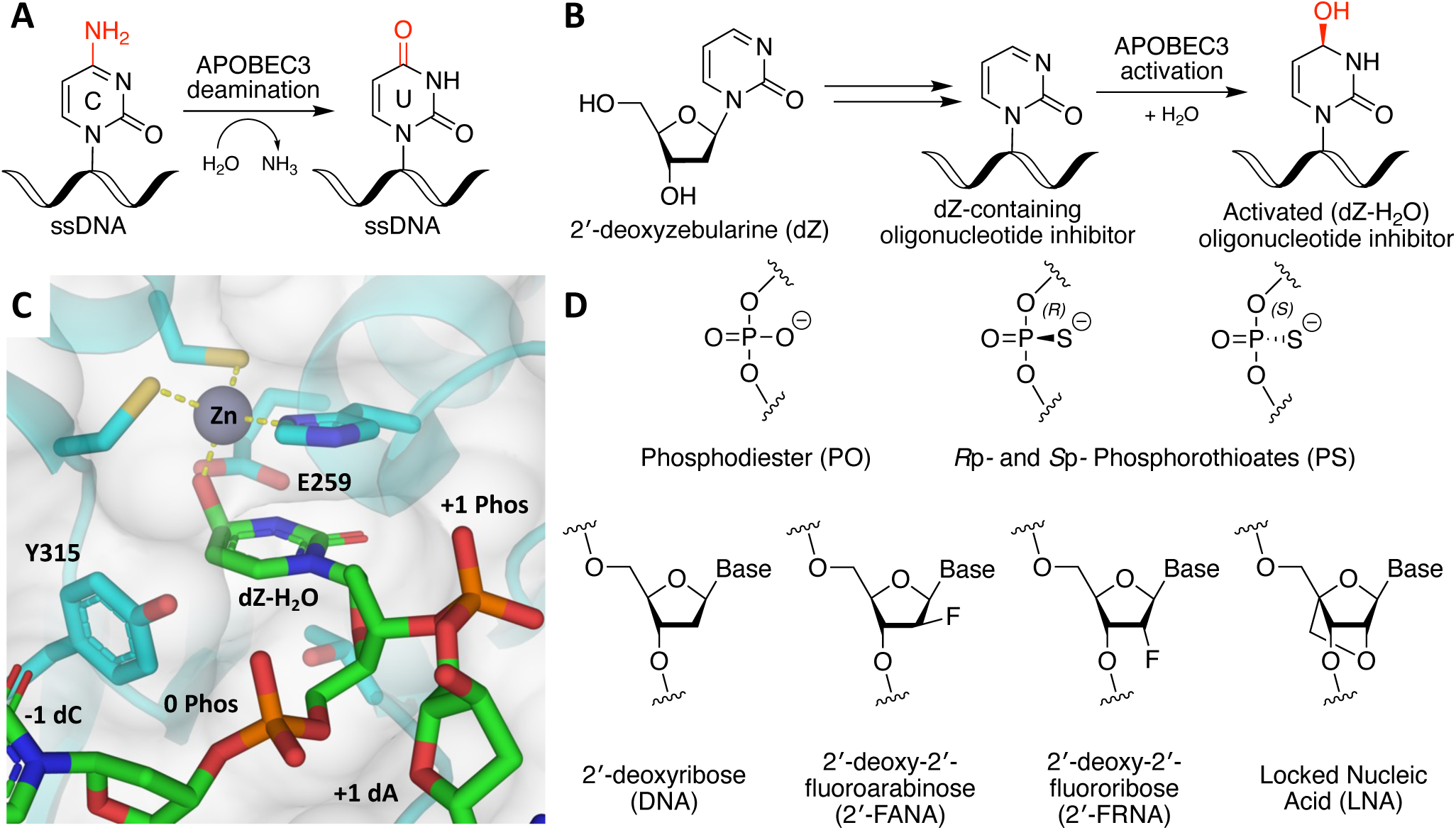
Structural basis of APOBEC3 activity and inhibition **A**) Diagram of APOBEC3 mediated deamination converting deoxycytidine to deoxyuridine in singlestranded DNA (ssDNA). **B**) Incorporation of 2′-deoxyzebularine (dZ) into short ssDNA oligonucleotides generates dZ-containing oligonucleotide inhibitors, which become activated by APOBEC3 enzymes forming a tighter-binding transition-state mimic (dZ-H2O). **C**) Recent co-crystal structure of a dZ-containing oligonucleotide inhibitor in complex with a APOBEC3G variant, confirming presence of the activated tetrahedral dZ-H2O bound in the active site (PDB:7UXD, Zn^2+^ is shown as a grey sphere, protein as cyan sticks/cartoon, and oligonucleotide inhibitor as green sticks). Central 3 nucleotides shown only. **D**) Chemical structures of common nucleic acid chemical modifications used in this work.

Despite this therapeutic interest, generating efficient small molecule inhibitors targeting A3 enzymes (15–18)has been difficult. Cytidine deaminase (CDA) shares a highly conserved active site with A3 enzymes (19–21); however, existing nucleoside based CDA inhibitors have no inhibitory potential against A3s (22), likely because A3s recognize longer single-stranded (ss) DNA substrates. Recent work to understand the substrate sequence specificities of A3s (23–25) and the determination of A3A (26,27) and A3G (28) co-crystal structures with substrate ssDNA have led to the design of substrate-mimicking oligonucleotide inhibitors. Commonly these inhibitors contain 2′-deoxy-zebularine (dZ), the deoxy analog of a potent CDA inhibitor, zebularine, in place of the target deoxycytidine (dC) in short ssDNA oligonucleotides. The dZ nucleobase is activated by the A3 enzyme, and follows the same deamination mechanism as dC, but becomes trapped in a tightly bound hydrated state that closely mimics the C to U transition state (**Fig 1B**). Recently we solved the first co-crystal structure of an oligonucleotide inhibitor bound to an active A3 enzyme, confirming the presence of an activated tetrahedral dZ-H_2_O nucleobase within the active site (**Fig 1C**) (19). Indeed, these dZ-containing 9to 13-nt oligonucleotides can inhibit A3A and A3G with potencies in the low μM to nM range (19,22,29,30) However, despite the promise of these A3 oligonucleotide inhibitors in terms of enzymatic potency, these molecules cannot be effective *in cellulo* and *in vivo* inhibitors. Unmodified single-stranded DNA is rapidly degraded by exo- and endonucleases, with unmodified sugar-phosphate backbones having a half-life on the order of minutes to hours within the cell (31,32). For any oligonucleotide-based compound to advance into cellular use, or even, one day, into the clinic, chemical modification of the sugars and/or phosphate groups are required (33,34). Fully chemically stabilized oligonucleotides (e.g antisense oligonucleotides and siRNAs) have been shown to have extended half-lives on the order of weeks to months (35). Arguably the most common modification to the phosphodiester (PO) backbone is the introduction of the phosphorothioate (PS) linkage (36) (**Fig 1D**), whereby one non-bridging phosphate oxygen is converted to sulfur, creating a chiral center (∼1:1 diastereomers) which is not controlled during conventional oligonucleotide synthesis. PS linkages greatly increase nuclease stability and binding to a variety of cellular proteins; this latter property drives increased cellular uptake but can also be associated with toxicity (37–40). Modifications can also be made to the sugar, most commonly at the 2′ position such as 2′-*O*-alkyl, 2′-fluoro, and 2′,4′-linked bicyclic derivatives (**Fig 1D**), and can increase nuclease stability as well as binding affinity to target nucleic acids (41–47).

To develop the next generation of A3 oligonucleotide inhibitors, we have combined our knowledge of the structural biology of A3-complexes and chemical modification of oligonucleotides to carry out a systematic and comprehensive evaluation of sugar- and phosphate-modified oligonucleotide inhibitors targeting both A3G and A3A. We show that fully PS-modified inhibitors have similar binding and inhibition characteristics to full PO inhibitors yet show drastically higher nuclease stability. However, due to the structural constraints of the active site of the A3 enzymes, we demonstrate that inhibitors lose potency if PS linkages are incorporated in the oligonucleotide immediately surrounding active site. Critically, mixed backbone oligonucleotide inhibitors retaining PO linkages flanking the target dZ but with PS linkages elsewhere can strongly increase potency while retaining much improved cellular stability. We also used molecular dynamics simulations to computationally guide the first reported incorporation of sugar modifications into A3G and A3A inhibitors. Most notably for A3G-CTD2, careful positioning of 2′-fluoro derivatives resulted in the most potent A3G-CTD2 inhibitor reported to date, with a *K*_i_ of 670 nM. Our best PS containing linear (single-stranded) inhibitors show increased potency compared to their unmodified PO counterparts, displaying single digit μM and low nM inhibition constants against A3G and A3A respectively. The potency and stability of our lead A3A targeting inhibitor was further enhanced by moving to a hairpin structure containing a mixed PO/PS backbone with modified sugars, resulting in the most potent A3A inhibitor reported to date, with a *K*_i_ of 9.2 nM. A panel of these inhibitors were tested in serum-nuclease assays, showing that only our chemically modified “second generation” inhibitors are stable enough to withstand nuclease degradation. These second generation, chemically modified, nuclease stable A3 inhibitors can now advance the field to move towards *in vitro* and *in vivo* applications, to further study the biological function of these enzymes and evaluate the therapeutic potential of A3 inhibition.

## METHODS

### Chemical Synthesis

General synthetic methods, scheme of synthetic route to dZ-phosphoramidite (**Supplementary Scheme 1**), and full NMR spectra and tabulations (^1^H, ^13^C, ^31^P) are given in supplementary data.

### 3′-5′-Di-O-(p-toluoyl)-2’-deoxyzebularine (1 in Supplementary Scheme S1)

2-Hydroxypyrimidine hydrochloride (10.21 g, 77.2 mmol, 2.5 eq) was suspended in hexamethyldisilazane (30 mL) with catalytic ammonium sulfate (0.204 g, 1.54 mmol, 0.02 eq) and refluxed at 140 °C for 2 hours under argon. The pale brown solution was then evaporated under reduced pressure, and used immediately in the following glycosylation reaction. For glycosylation, this crude silylated 2-hydroxypyrimidine was resuspended in chloroform (250 mL) in a three-neck flask fitted with a distillation condenser. This solution was stirred and vigorously heated at 100 °C with continuous addition of chloroform (∼2-3 mL/min) to maintain constant volume. Hoffer’s chlorosugar (12.00 g, 30.9 mmol, 1.0 eq) was dissolved in chloroform (80 mL) and added to the boiling solution over 30 minutes, during which time the trimethylsilylchloride byproduct was distilled off. Once addition was complete, heating was maintained for a further 10 minutes before cooling to RT, and the reaction mixture was extracted with 250 mL each of water, then saturated NaCl, and dried over Na_2_SO_4_. Volatiles were removed under reduced pressure to give quantitative yield of a white solid, of good purity and 2:1 β -selectivity by NMR. TLC: 10 % MeOH in CH_2_Cl_2_; *R_f_* = 0.72. The α-anomer was removed selectively based on its insolubility in 40 % choroform/hexane at -20 °C overnight, and three successive cycles gave (9.80 g, 71 %) with a final β /α ratio of 6.3:1.

### 2′-deoxyzebularine (2 in Supplementary Scheme S1)

Toluoyl-protected 2’-deoxyzebularine (**1**) (9.8 g, 21.9 mmol) was dissolved in a minimum amount of anhydrous dichloromethane, transferred to a glass pressure vessel, and combined with 7N methanolic ammonia (400 mL). After stirring at RT for 28 hours, the volatiles were removed under reduced pressure, and the residue dissolved in water (250 mL), and extracted 5x with chloroform (150 mL). The aqueous portion was then lyophilized to give a white solid (4.23 g, 91 %) of good purity and 7.3:1 β -selectivity by NMR. TLC: 10 % MeOH in CH_2_Cl_2_; *R_f_* = 0.15.

### 5′-O-(4,4′-dimethoxytrityl)-2′-deoxyzebularine (3 in Supplementary Scheme S1)

Nucleoside (**2**) (3.58 g, 16.9 mmol, 1.0 eq) was dissolved in anhydrous pyridine (170 mL) and cooled on ice under argon. 4,4′-Dimethoxytrityl chloride (6.87 g, 20.3 mmol, 1.2 eq) was added and the reaction allowed to warm to RT and stirred for 16 hours. The reaction mixture was concentrated to an orange gum under reduced pressure, dissolved in dichloromethane (250 mL), and extracted with 150 mL each water, saturated bicarbonate, and saturated brine, and then dried over Na_2_SO_4_. The residue was taken up in dichloromethane and flash purified using an isocratic gradient of 0 %, then 0-6 % methanol in dichloromethane. The column was pre-neutralized with dichloromethane containing 1 % triethylamine. Fractions containing the pure β -anomer (which eluted first) were pooled, evaporated under reduced pressure, and coevaporated with chloroform to give a crunchy white solid (6.17 g, 71 % based on total input mass of (**2**), or 82 % in terms of β -anomer). TLC: 5 % MeOH in EtOAc; *R_f_* = 0.45

### 5′-O-(4,4′-dimethoxytrityl)-2′-deoxyzebularine,3’-[(2-cyanoethyl)-(N,N-diisopropyl)]-phosphoramidite (4 in Supplementary Scheme S1)

Tritylated nucleoside (**3**) (1.78 g, 3.46 mmol, 1.0 eq) was dissolved in anhydrous dichloromethane (20 mL), cooled to 0 °C on ice, and anhydrous diisopropylethylamine (1.69 mL, 9.68 mmol, 2.8 eq) was added. Under argon, 2-cyanoethyl-*N*,*N*-diisopropylchlorophosphoramidite (1.07 mL, 4.84 mmol, 1.4 eq) was then added and the solution stirred on ice for 2 hours, until TLC indicated reaction completion. The reaction mixture was extracted with saturated bicarbonate (20 mL), the organic phase dried over Na_2_SO_4_, and evaporated under reduced pressure. The residue was taken up in dichloromethane and flash purified using an isocratic gradient of 0 %, then 0-4 % methanol in dichloromethane. The column was pre-neutralized with dichloromethane containing 1 % triethylamine. Fractions containing the pure phosphoramidite were pooled, evaporated under reduced pressure, and coevaporated with chloroform to give a crunchy off-white foam (1.99 g, 80 %), as ∼1:1 phosphoramidite diastereomers at phosphorus by ^31^P NMR. TLC: 5 % MeOH in EtOAc; *R_f_* = 0.72

### Oligonucleotide Synthesis, Purification, Characterization

Oligonucleotide substrates were purchased from IDT with standard desalting and reconstituted in nuclease-free water for use. All dZ-containing oligonucleotides were synthesized in-house using an either an Akta OligoPilot or DrOligo48 synthesizer using standard methods, and all reagents were purchased from ChemGenes (Wilmington, MA). All oligonucleotides were synthesized using 1000 Å long-chain alkyl amine (LCAA) controlled pore glass (CPG) functionalized with the first nucleotide. Phosphoramidites were purchased from ChemGenes except the dZ phosphoramidite which was synthesized as above, and were diluted to 0.1 M in anhydrous acetonitrile (ACN). Detritylation was achieved using 3% trichloroacetic acid (TCA) in toluene or dichloromethane. Oxidation was accomplished using 0.05 M iodine in water/pyridine (9:1 v/v), and sulfurization using 0.1 M 3-[(dimethylaminomethylene)amino]-3H-1,2,4dithiazole-5-thione (DDTT) in pyridine. 0.25 M 5-(Benzylthio)-1H-tetrazole (BTT) in ACN was used as the activator. Capping was achieved by Cap A (20% n-methylimidazole in ACN), and Cap B (20% acetic anhydride, 30% 2,6-lutidine in ACN), however capping was skipped for hairpin A3A inhibitors to prevent acetylation of dG-PAC. Backbone cyanoethyl deprotection was carried out on-support using 10% diethylamine in ACN.

Oligonucleotides were cleaved from the solid support and deprotected by treatment with conc. aqueous ammonia at RT for 4-6 hours, then vacuum concentrated, resuspended in 5 % ACN, filtered, and purified by semi-preparative HPLC.

All inhibitors were purified by ion-exchange HPLC using gradients of up to 0.5M NaBr in 10% aqueous ACN with 20 mM NaOAc. Hairpin inhibitors were first purified by ion-paired reverse-phase (IPRP) HPLC using gradients of 5 to 50% methanol in an aqueous solution containing 400 mM hexafluoroisopropanol (HFIP) and 15 mM triethylamine (TEA). LC peaks were monitored at 260 nm, and the major peak was fractionated and analyzed by LC-MS. Pure fractions were pooled and concentrated to dryness by vacuum centrifugation.

Oligonucleotides were then re-suspended in 5% ACN and desalted by size exclusion chromatography on a 25 × 250 mm custom column packed with Sephadex G-25 media (Cytiva, MA) and lyophilized. Final desalted oligonucleotides were re-suspended in nuclease free water at stock concentrations of 1-2 mM and stored at -20 °C until use. Oligonucleotide extinction coefficients were calculated using published values for monomers, using a value of 500 Lmol^-1^cm^-1^ at 260 nm for dZ.

All oligonucleotides were characterized by LC-MS analysis on an Agilent 6530 accurate mass Q-TOF using an AdvanceBio C18 oligonucleotide column (Agilent), at 60 °C, eluting with a gradient of increasing methanol in 100 mM HFIP and 9mM TEA in LC-MS grade water. MS parameters: Source, electrospray ionization; ion polarity, negative mode; range, 100–3,200 m/z; scan rate, 2 spectra/s; capillary voltage, 4,000; fragmentor, 180 V.

### Molecular Modelling

The cocrystal structure of A3G-CTD2 bound to ssDNA (PDB ID: 6BUX(28)) was used for molecular modeling of A3G with inhibitors. The active site cytidine was converted to the catalytically hydrated tetrahedral dZ-H_2_O; 4-(R)-hydroxy-3,4-dihydro-2′-deoxy-zebularine and A259 was mutated to Glu to simulate the active form of the enzyme, using Maestro 3D Builder(Schrödinger). We note this active form of dZ-H_2_O we have since confirmed in a cocrystal structure (PDB ID: 7UXD)(19).

The cocrystal structure of A3A bound to ssDNA (PDB ID: 5SWW(27)) was used for molecular modeling of A3A with inhibitors. The DNA sequence was mutated in Coot(48) and Maestro 3D Builder to match that of the central region of the linear inhibitor sequence, and the active site cytidine was converted to the catalytically hydrated tetrahedral dZ-H_2_O; 4-(R)-hydroxy-3,4-dihydro-2′-deoxy-zebularine. A72 was mutated to Glu to simulate the active form of the enzyme, and residues 44-47 (amino acids TSVK), were built into the missing electron density using Maestro 3D Builder.

All additional chemical modifications to the oligonucleotide inhibitors were performed using Maestro 3D Builder.

### Molecular Dynamics Simulations and Analysis

Molecular dynamics simulations were performed using Desmond (Schrödinger). Models were first optimized using Protein Preparation Wizard at pH 6.5. Simulation systems were built using Desmond System Setup, utilizing the SPC solvation model, cubic boundary conditions of 10 Å buffer box size, and OPLS3 force field. The final simulation systems were neutral and had 0.15M NaCl. A multistage MD simulation protocol previously described(49) was used to simulate triplicates for 200 ns each.

Simulation trajectory RMSDs were calculated using the RMSD Visualizer Tool in VMD. Hydrogen bond occupancies were calculated using ROBUST(50) with a cutoff of 3.0 Å, a donor angle of at least 120°, and an acceptor angle of at least 90°. Visual studies and analysis of trajectories was performed in VMD(51) and PyMol(52).

### Protein Expression and Purification of A3G-CTD2

A3G-CTD2 was expressed and purified from *E. coli* BL21 cells as previously described(19) with some minor modifications (see Supplementary Information for full details). Throughout the purification, sample purity was accessed by SDS-PAGE. A3G-CTD2 was quantified (molar extinction 40,450 M^-1^ cm^-1^), aliquoted, and stored at -80 °C in 50 mM sodium phosphate pH 7.3, 150 mM NaCl, 50 μM zinc chloride, 0.002 % Tween-20, 10 % glycerol and 1 mM DTT.

### Cell-Free Protein Expression and Purification of WT-A3A

Due to the cytotoxic nature of A3A, a cell free protein expression system based on wheat germ extract was used to express protein for *in vitro* studies. The C-terminal 6xHis tagged A3A gene was cloned into CellFree Sciences’ (Matsuyama, Ehime, Japan) expression vector pEU-E01-MCS using the 5’ Xhol and 3’ Notl cloning sites. This plasmid was used in conjunction with CellFree Sciences’ WEPRO®7240H Core Kit to express the A3A protein at a 6 mL expression reaction scale following manufacturer’s directions. In brief, RNA transcription was setup in 250 μL in Transcription Reaction Buffer LM, 2.5 mM NTP mix, 1 U/μL RNase inhibitor, 1 U/μL SP6 RNA Polymerase, and 100 ng/μL APOBEC3A plasmid. Reaction was incubated for 6 hours at 37 °C and protein expression was initiated by adding 250 μL of WEPRO®7240H (240 OD), 40 ng/mL Creatine Kinase, 10 μM Zn acetate in SUB-AMIX® SGC reaction buffer for 20 hours at 15 °C. The sample was mixed with binding buffer (25 mM Tris pH 7.5, 250 mM NaCl, 1 mM TCEP, 0.002 % TWEEN20, 10 μM ZnCl_2_, and 50 mM Imidazole) and incubated with 0.2 mL of equilibrated Ni Sepharose resin for 1 hour at 4 °C. Protein-bound resin was washed 2 times with 2 mL of buffer containing 25 mM Tris pH 7.5, 500 mM NaCl, 1 mM TCEP, 0.002 % TWEEN20, and 10 μM ZnCl_2_ and 2 times with 2mL of buffer with 25 mM Tris pH 7.5, 250 mM NaCl, 1 mM TCEP, 0.002 % TWEEN20, and 10 μM ZnCl_2_. Protein was eluted with buffer containing 25 mM Tris pH 7.5, 250 mM NaCl, 1 mM TCEP, 0.002 % TWEEN20, 10 μM ZnCl_2_, and 400 mM imidazole. The eluted protein was buffer exchanged into 50 mM sodium phosphate, pH 7.5, 100 mM NaCl, 1 mM DTT, and 0.002 % Tween-20 using Zeba protein desalting columns with 7K MWCO resin. Throughout the purification, sample purity was accessed by SDS-PAGE.

### NMR Deamination Assay

Initial rates of cytosine deamination with and without inhibitor were measured at 25 °C using a ^1^H-based NMR deamination assay, in which formation of the uracil product is measured by integration of the unique H5 doublet over time, as previously described (28). All experiments were performed with 200 µM substrate, and in buffer (50 mM NaPO4 pH 6.5, 100 mM NaCl, 1 mM DTT, 10 μM ZnCl_2_, and 0.002 % Tween 20) with 10 % D_2_O.

For A3G-CTD2, NMR data was collected on a Bruker Ascend 600 MHz NMR spectrometer equipped with cryoprobe. A3G-CTD2 (50 nM) was incubated with substrate (5′-AATCCCAAA), and inhibitors were screened at 5 μM, and *K*_i_’s (see *K*_i_ Determination) were determined using 0-20 μM inhibitor. All experiments were conducted in duplicate. Data was analyzed in MestReNova v14, and product formation calculated by integrating the H5 uracil doublet at 5.66 ppm relative to a singlet at 8.16 ppm.

For A3A, NMR data was collected on a Bruker Avance III 600 MHz NMR spectrometer equipped with cryoprobe. A3A (50 nM for linear inhibitors, 10 nM for hairpins) was incubated with substrate (5′TTCAT), inhibitors were screened at 5 μM (linear) or 100 nM (hairpins), and *K*_i_’s (see *K*_i_ Determination) were determined using 0-5 μM (linear) or 0-10 nM (hairpin) inhibitor. All experiments were conducted in triplicate. Data was analyzed in Topspin 4.1, and product formation calculated by integrating the H5 uracil doublet at 5.71 ppm relative to external standard.

### ***K*_i_** determination

For *K*_i_ determination, all compounds were treated as competitive inhibitors. Slopes were obtained from linear regression of initial rates (V) of product formation over time, under varying inhibitor concentrations. From here, 1/V vs [I] was plotted to give a Dixon plot. A linear fit (giving equation (y = ax + b)) of the Dixon plot provides a slope (*a*) and intercept (*b*) containing *K*_m_, kcat and *K*_i_. *K*_i_ is determined from 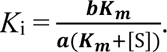 All errors were calculated using error propagation.

We previously measured the *K*_m_ value for this A3G substrate (5′-AATCCCAAA) as, *K*_m_: 0.509 ± 0.086 mM (19)). For A3A, we determined the *K*_m_ for the 5′-TTCAT substrate *K*_m_: 226 ± 19 µM, using 0.1-1 mM substrate by NMR deamination assay, described above. 200 µM substrate ([S]) was used for all inhibition experiments.

### Expression and Purification of inactive A3A for MST

A3A-E72A inactive mutant was expressed from pGEX6P-1 expression plasmid with an N-terminal Glutathione S-transferase (GST) tag and a C-terminal 6×His tag. Expression plasmid was transformed into chemically competent BL21(DE3) cells and selected by Ampicillin. Selected cells were grown at 37°C in LB media with Ampicillin to 0.6 OD600. The culture was chilled to 17°C for 1 hour and protein expression was induced for 18 hours with 0.2 mM isopropyl β-D-1-thiogalactopyranoside (IPTG). All the steps for protein purification were performed at 4 °C. *E. coli* cells were harvested by centrifugation and re-suspended in lysis buffer (50 mM sodium phosphate pH 7.5, 150 mM NaCl, 2 mM DTT, and 0.002% Tween-20) and EDTA free protease inhibitor cocktail (Roche, Basel, Switzerland). The suspended cells were disrupted using Avestin C3 homogenizer. Cell debris was separated by centrifugation at 48,000×g for 30 minutes. Supernatant containing desired protein was applied to Glutathione-Sepharose resin (GE Healthcare Life Science) equilibrated with lysis buffer and agitated for 2 hours. Protein-bound resin was washed with Pre-Scission Protease cleavage buffer (50 mM sodium phosphate, pH 7.5, 100 mM NaCl, 2 mM DTT, and 0.002% Tween-20) and incubated with Pre-Scission protease (GE Healthcare Life Science) for 18 hours. The supernatant containing the cleaved protein was separated from the resin by centrifugation and further purified using a HiLoad 16/600 Superdex 75 gel filtration column (GE Healthcare Life Science) equilibrated with 50 mM sodium phosphate pH 7.5, 100 mM NaCl, 1 mM DTT, and 0.002% Tween-20. Protein purity was analyzed by SDS-PAGE.

### Microscale Thermophoresis measurements

The affinity of WT-A3A to dZ-inhibitors and A3A-E72A to substrates were determined as dissociation constants (*K*d) using a Monolith (NanoTemper Technologies, GmbH, Munich, Germany) microscale thermophoresis instrument. RED-tris-NTA fluorescent dye solution was prepared at 100 nM in the MST buffer (50 mM sodium phosphate pH 7.5, 100 mM NaCl, 1 mM DTT, 0.002 % Tween 20). A3A was mixed with dye at 100 nM and incubated for 30 min at room temperature followed by centrifugation at 15,000 × g for 10 min. Dye labeled A3A solutions were mixed with inhibitor for final protein concentration of 50 nM and final inhibitor concentrations of 100 µM to 3 nM for linear inhibitors and 5 µM to 0.15 nM for hairpin inhibitors with 2-fold dilutions. Protein-inhibitor solutions were incubated at 25 °C for 1 hour before MST measurements using NanoTemper MST premium capillaries. Measurements were performed at 25 °C with 100 % excitation power and 40 % MST power. The experiment was repeated three times and data analysis was carried out using MO affinity analysis software (NanoTemper Technologies). Data presented as relative fold change in main figures, primary binding data with uncertainties given in Supplementary Tables S2 and S3.

### Serum nuclease HPLC assay

For each experiment, 18 nmol of oligonucleotide in 54 μL Dulbecco’s Modified Eagle Medium (DMEM) (Sigma, #F0926) was mixed with 216 μL of 25 % Fetal bovine serum (ThermoFisher, #11995065) in DMEM, and incubated at 37 °C in capped eppendorfs (20 % final FBS). Aliquots were taken at various timepoints over 72 hours by transferring 30 uL of the reaction mixture (2 nmol inhibitor) into 20 uL of formamide to quench the reaction, immediately flash frozen in liquid nitrogen, then stored at -80 °C until HPLC analysis using a 1260 Infinity HPLC (Agilent). Example HPLC trace shown in **Supplementary Fig S3**.

All inhibitors except H1 were analyzed by ion-exchange HPLC using gradients of up to 0.5M NaBr in 10% aqueous ACN with 20 mM NaOAc. Due to poor resolution, H1 was analyzed using reverse-phase HPLC using gradients of 2 to 35 % ACN in an aqueous solution containing 100 mM hexylammonium acetate (HAA). The relative A260 peak area of the full-length peak was calculated as a percentage of the total area of all peaks eluting over the gradient phase, and normalized to the peak area at time zero. Experiments were conducted in duplicate. Data was plotted and analyzed in GraphPad Prism10.

## RESULTS

### Improved synthesis of dZ-phosphoramidite

To synthesize our oligonucleotide-based inhibitors, we first synthesized the 2′-deoxy-zebularine (dZ) phosphoramidite based on a previous method by Kvach *et al.* (22), with improvements to both synthetic yield and simplicity (**Supplementary Scheme S1** and methods). Briefly, 2-hydroxypyrimidine hydrochloride was silylated by refluxing in hexamethyldisilazane with catalytic ammonium sulphate. This solution of silylated nucleobase was then evaporated under reduced pressure, resuspended in chloroform, and used immediately in the following glycosylation reaction. Hoffer′s chlorosugar was added dropwise to the silylated nucleobase in vigorously boiling chloroform under distillation to remove the trimethylsilylchloride byproduct; previously shown to increase β -anomer selectivity(53). Extraction and evaporation gave the crude protected dZ nucleoside in quantitative yield and good purity, with 2:1 β -selectivity. The α-anomer was removed on the basis of its insolubility in 40% chloroform in hexane, at -20 °C to give a 6.3:1 anomeric ratio. Treatment with 7N ammonia in methanol overnight furnished the deprotected dZ nucleoside. This was 5′-dimethoxytrityl (DMT) protected and flash purified to give exclusively the β -anomer, which was then phosphitylated to provide the final dZ-phosphoramidite, as an equal mixture of diastereomers at phosphorus. By avoiding vacuum distillation of the silylated-nucleobase and minimizing chromatography steps, our method resulted in a ∼50 % increase in final dZ-phosphoramidite yield compared to previous literature(22).

### Synthesis of chemically modified dZ-containing oligonucleotide inhibitors

Oligonucleotides were synthesized using standard phosphoramidite chemistry (see methods). However, we found that the dZ unit was sensitive to base treatment during the oligonucleotide cleavage/deprotection step and required mild conditions. Namely, treatment of the controlled pore glass (CPG) support with standard deprotection conditions (concentrated aqueous NH3 for 16 hours at 55 °C, or 1:1 concentrated aqueous NH3:methylamine (AMA) for 1.5 hours at room temperature or 10 minutes at 65 °C) gave complete destruction by mass-spectrometry, with formation of -36 and -78 Da products whose structures we could not determine (**Supplementary Fig S1**). Incubation with concentrated aqueous NH3 for 16 hours at room temperature showed minor degradation, while 6 to 8 hours at room temperature was optimal, allowing full deprotection of benzoyl and dimethylformamidine groups (from A and G, respectively) without damage to dZ. Therefore, isobutyryl protection and universal controlled pore glass (CPG) supports must be avoided for the synthesis of dZ-containing oligonucleotides, due to the harsher deprotection conditions they require.

Oligonucleotide sequences were based on previously characterized length and sequence preferences for A3G and A3A substrates, replacing the target dC with dZ to form inhibitors. For A3G we used inhibitors containing the preferred 5′-CC<u>C</u>Abinding motif(24,25): (5′-AATCC<u>dZ</u>AAA). For A3A we used inhibitors containing the preferred 5′-(T/C)T<u>C</u>(A/G)binding motif(23): (5′-AAATT<u>dZ</u>AAAAAAA or 5′-TGCGCTT<u>dZ</u>GCGCA for hairpins). Oligonucleotides are numbered with a format (e.g **I1G**, **S1A**, or **H3**_**A**_) where the prefix denotes **<u>S</u>**ubstrate containing dC, **<u>I</u>**nhibitor containing dZ, or a **<u>H</u>**airpin-structured dZ inhibitor, and subscript denotes an A3**<u>A</u>** or A3**<u>G</u>** targeting sequence. A full list of synthesized inhibitors and their characterization is shown in **Supplementary Table S1**. These inhibitor sequences match the substrate specificity of each enzyme, allowing us to carefully characterize the impact of chemical modifications on both A3G and A3A.

### Phosphorothioate modification pattern impacts A3G inhibition

To evaluate the effect of nuclease stable phosphorothioate (PS) linkages on the inhibition of A3G, we tested a panel of PS modified oligonucleotide inhibitors (**Fig 2**) against a catalytically active and soluble A3G-C-terminal-domain variant (referred to throughout as A3G-CTD2), using a previously validated ^1^H-based NMR deamination assay(54) (28). We characterized the unmodified full PO A3G inhibitor **I1G**, as a reference and determined a *K*i of 3.34 ± 0.70 μM, consistent with our previous work(19). We then tested a panel of modified inhibitors containing PS linkages either across the entire sequence or restricting the modification to either adjacent or remote from the target dZ (**Fig 2A, B**). These inhibitors were first screened at 5 μM (**Fig 2B**) with 50 nM A3G-CTD, where **I1G** inhibited 49% of A3G-CTD2’s activity. Inhibitors with PS modifications within the active site, including the fully modified **I2G** as well as **I3G** and **I6G**, all lost inhibition relative to **I1G**. Interestingly, introducing PS modifications away from the linkages surrounding target dZ, as in **I4G** and **I5G**, enhanced potency compared to **I1G**.

**Figure 2.**
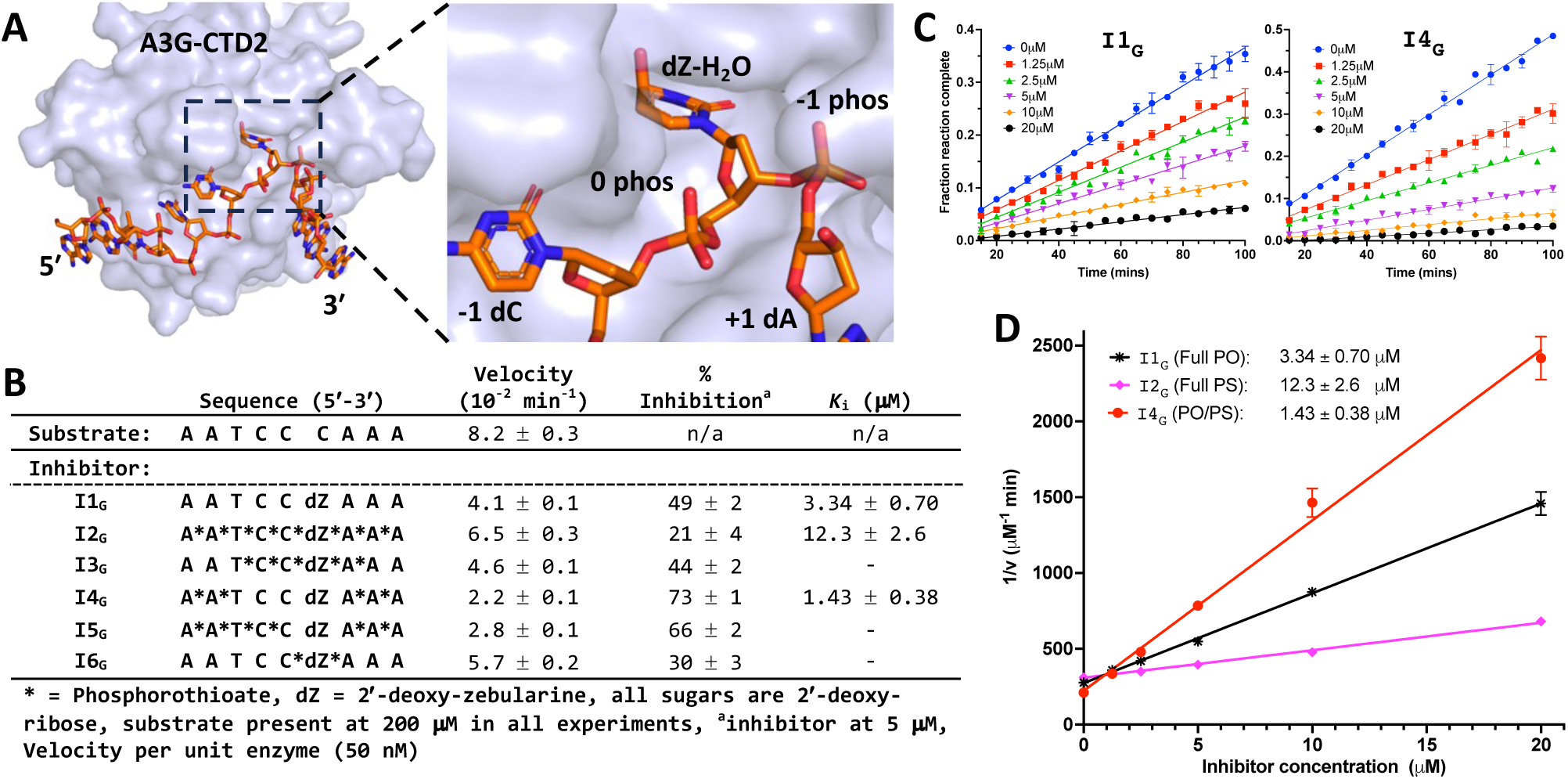
Evaluation of phosphorothioate modification pattern on A3G inhibition **A**) Structural model of inhibitor **I1G** bound to A3G-CTD2. Left; global view of the 9-nt oligonucleotide bound across the enzyme. Right; active site region showing activated dZ-H2O and flanking phosphodiester linkages. Model based on 6BUX structure with A259E, and active site dC changed to dZ-H2O. (A3G-CTD2; lilac, surface representation. **I1G** inhibitor; orange, sticks). **B**) Table of phosphorothioate modified oligonucleotides and inhibition parameters against A3G-CTD2 measured by ^1^H NMR deamination assay, percentage inhibition is relative to substrate only deamination reaction velocity. **C**) Representative dose dependent inhibition plots for **I1g** and **I4g** inhibitors, across multiple concentrations (datapoints duplicate, error bars SEM). **D**) Inverse velocity vs inhibitor concentration plot (Dixon plot) used to determine inhibition constants (*K*i) from the slope (see SI). Steeper line slopes indicate stronger inhibitor (datapoints duplicate, error bars SEM).

Based on these results, we carried out dose-dependent inhibition experiments (**Fig 2C**) to determine *K*i values (**Fig 2D**) for the most potent and most heavily modified inhibitors, compared to **I1G**. Fully PS modified **I2G** showed a 3.7-fold loss in potency with a *K*i of 12.2 ± 2.6 μM. Inhibitor **I4G**, with two PS linkages on each end of the oligonucleotide (four in total) was the most potent with a *K*i of 1.43 ± 0.38 μM, 2.3-fold more potent than the full PO inhibitor **I1G**. Thus, likely due to the structural constraints around the active site of A3G, modification of the phosphates close to the target dZ with diastereomeric PS linkages weakens inhibitor binding, while introduction of PS linkages distal from the active site enhances A3G inhibition.

### Phosphorothioate modification pattern also impacts A3A inhibition

To assess whether similar trends occur with the more therapeutically relevant A3A enzyme, we next investigated the impact of PS modification pattern on A3A binding and inhibition (**Fig 3**). Using an A3Apreferred oligonucleotide substrate sequence, we tested a similar panel of PS modification patterns in sequence-matched substrates or inhibitors, varying the extent of modification around the A3A active site (**Fig 3A-C**). This allowed the testing of oligonucleotide binding affinity to both the active enzyme (for dZ-containing inhibitors) and the catalytically inactive variant A3A-E72A (for dC-containing substrates), using microscale thermophoresis (MST) (**Fig 3B**). Overall, there was a good correlation between binding affinity measured by MST, and inhibition of deaminase activity. As with A3G, the ^1^Hbased NMR deamination assay was used to screen inhibitors at 5 μM (**Fig 3C**), and to determine inhibition constants for the most potent and most heavily modified sequences (**Fig 3C-E**). For the unmodified, full PO A3A inhibitor **I1**_**A**_ (64% inhibition at 5μM), we measured a *K*i of 342 ± 62 nM. The fully PS modified A3A inhibitor **I2**_**A**_, and its equivalent dC-containing substrate **S2A**, both showed a slight loss of binding affinity, and a 1.4-fold decrease in inhibition potency with a *K*i of 469 ± 56 nM. However, A3A was particularly sensitive to PS modifications at two linkages directly flanking 5′ and 3′ of the target dZ (or dC). Addition of PS at only these two positions resulted in by far the weakest binding substrate or inhibitor, with a 4to 13-fold loss in binding affinity (**S4A** and **I4**_**A**_ respectively, **Fig 3B**), and the poorest inhibition (22% by **I4**_**A**_, **Fig 3C**). Remarkably, removal of PS from these two flanking positions from an otherwise fully PS modified sequence, gave the tightest binding substrate or inhibitor (**S3A** or **I3A, Fig 3B**), and resulted in the most potent A3A inhibitor **I3**_**A**_ with a *K*i of 51 ± 11 nM (**Fig 3C-E**). These results demonstrate that A3A, like A3G-CTD2, highly disfavors diastereomeric PS linkages flanking the active site. However, A3A binding and inhibition is significantly improved for substrates and inhibitors containing PS linkages at all but the flanking linkages either side of the target dZ, likely due to the tight constraints of the active site structural architecture.

**Figure 3.**
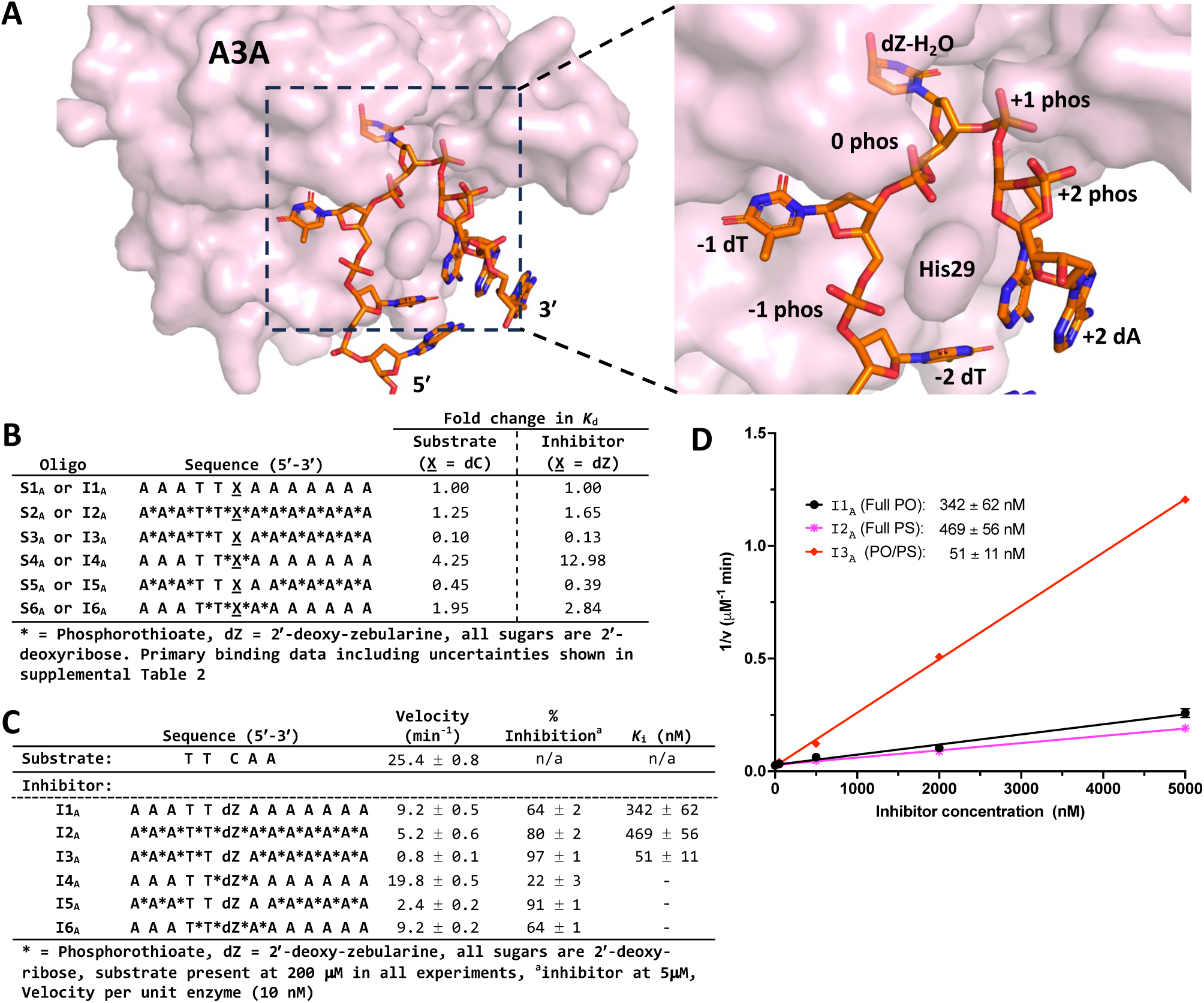
Evaluation of phosphorothioate modification pattern on A3A inhibition **A)** Structural model of the central portion of inhibitor **I1^A^** bound to WT-A3A. Left; global view of the central 7-nt of the oligonucleotide bound in a U-shape conformation. Right, active site region showing activated dZ-H^2^O, and flanking phosphodiester linkages, with the DNA wrapped around the central His29. Model based on 5SWW structure with A72E, active site dC changed to dZ-H^2^O, nucleobases changed from ATCGGG to ATTdZAAA. (A3A; pink, surface representation. **I1^A^** inhibitor; orange, sticks). **B)** Table of phosphorothioate modified substrates and inhibitors and their relative binding affinities (*K*^d^) to A3A measured by MST, data shown as fold change relative to full PO control. **C)** Table of phosphorothioate modified oligonucleotides and inhibition parameters against A3A measured by ^1^H NMR deamination assay, percentage inhibition is relative to substrate only deamination reaction velocity. **D)** Inverse velocity vs inhibitor plot (Dixon plot) used to determine inhibition constants (*K*^i^) from the slope (see SI). Steeper line slopes indicate a stronger inhibitor (datapoints triplicate, error bars SD).

### A3 enzymes are sensitive to placement of sugar modifications within oligonucleotide inhibitors

To further investigate strategies to optimize the stability and potency of our oligonucleotide inhibitors we next incorporated a selection of sugar modifications into both A3G and A3A inhibitors (**Fig 4**). We focused on 3 common modifications, 2′-deoxy-2′-fluoroarabinonucleic acid (2′-FANA), 2′-deoxy-2′fluororibonucleic acid (2′-FRNA), and 2′-*O*,4′-*C*-methylene-*β*-D-ribonucleic acid (LNA) (**Fig 1)**. We first examined the co-crystal structure of A3G-CTD2 bound to a ssDNA substrate (6BUX) to determine the sugar conformation at each DNA nucleotide (C2′- or C3′-endo). We then designed two initial inhibitors containing 2′-FANA or LNA sugar modifications, introduced at positions where the modification was predicted to reinforce the observed sugar conformation, termed **I7G** and **I8G**. To our surprise, these inhibitors lost all potency relative to **I1G** at 5 μM (3% and 5% vs 49%), and showed negligible inhibitory activity even at 50 μM (**Supplementary Fig S2**).

**Figure 4.**
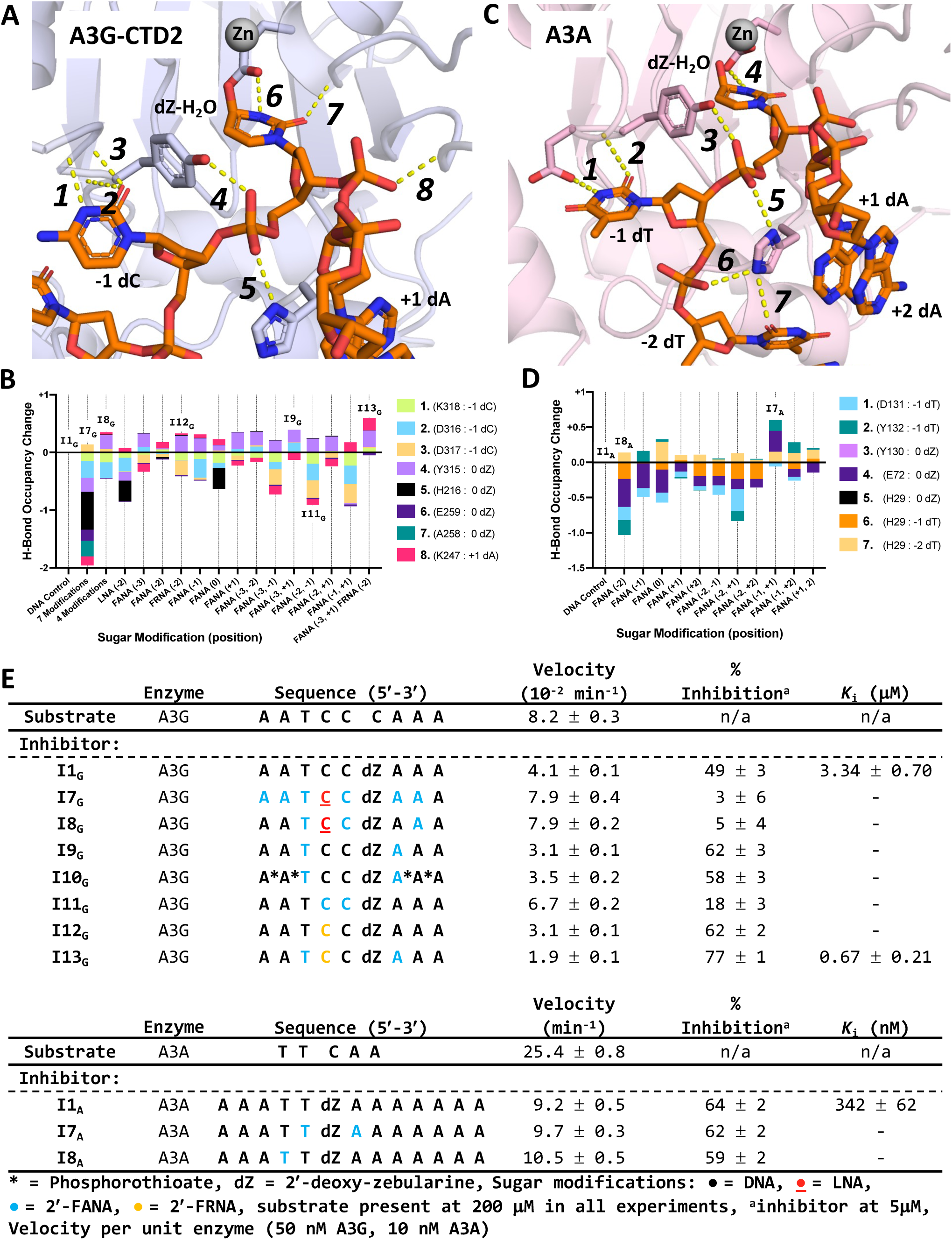
Evaluation of sugar modification pattern on A3G and A3A inhibition **A)** Structural model of the central portion of inhibitor **I1^G^** bound to A3G-CTD2 showing 8 predicted key protein-inhibitor hydrogen bonds. Model based on 6BUX structure with A259E, and active site dC changed to dZ-H^2^O. (A3G-CTD2; lilac, cartoon/sticks. **I1^G^** inhibitor; orange, sticks. Key hydrogen bonds; dashed yellow lines; numbers correspond to interaction in panel **B**). **B**) Plot of cumulative hydrogen bond occupancy change vs DNA only reference **I1^G^**, predicted by molecular dynamics simulation, for a range of sugar-modified A3G-CTD2 inhibitors). Legend (right) corresponds to interactions shown in **A**. **C)** Structural model of the central portion of inhibitor **I1^A^** bound to A3A showing 7 predicted key protein-inhibitor hydrogen bonds. Model based on 5SWW structure with A72E, active site dC changed to dZH^2^O, nucleobases changed from ATCGGG to ATTdZAAA. (A3A; pink, cartoon/sticks. **I1^A^** inhibitor; orange, sticks) Key hydrogen bonds; dashed yellow lines; numbers correspond to interaction in panel **D**). **D**) Plot of cumulative hydrogen bond occupancy change vs DNA only reference **I1^A^**, predicted by molecular dynamics simulation, for a range of sugar-modified A3A inhibitors). Legend (right) corresponds to interactions shown in **C**. **E)** Table of sugar modified inhibitors and inhibition parameters against A3GCTD2 (top) and A3A (bottom) measured by ^1^H NMR deamination assay, percentage inhibition is relative to substrate only deamination reaction velocity.

As optimal oligonucleotide binding involves a large combination of sugar conformations and possible modifications, we turned to molecular modeling to guide modification choice and placement. Starting from DNA-bound co-crystal structures of A3G-CTD2 (6BUX) (28) and A3A (5SWW) (27) we changed the DNA sequence to match our respective inhibitors, and modelled in sugar modifications at different positions, resulting in 27 various designs. We performed 200-nanosecond parallel molecular dynamics simulations (pMD)(55,56) in triplicate for each complex, tracking changes to key hydrogen bonds between each oligonucleotide inhibitor and the respective enzyme (**Fig 4A, B**). Changes in hydrogen bonding occupancies compared to the initial DNA-only reference inhibitors (**I1G** or **I1**_**A**_) were plotted for all design options (**Fig 4C, D**). Based on these results, we chose an additional 7 sugar-modified inhibitors for synthesis and characterization against A3G and A3A (**Fig 4E**).

For A3A, we synthesized two sugar-modified inhibitors, one predicted to stabilize hydrogen bonding interactions and the other predicted to reduce them. However, both compounds showed similar inhibition relative to the non-sugar-modified control **I1**_**A**_. Therefore, for A3A our analysis of computationally predicted hydrogen bond occupancies was insufficient to meaningfully inform further increases in inhibitory potency.

For A3G-CTD2, the approach was more successful. The heavily modified **I7G** was predicted to show a considerable loss in hydrogen bond occupancy, consistent with its poor experimental inhibition as described above. Similarly, **I11G** showed agreement between computationally predicted and experimentally measured loss of inhibition. Interestingly, inhibitor **I9G** with 2′-FANA incorporated at the -3 and +1 positions was predicted to show enhanced overall hydrogen bonding occupancy compared to **I1G**, and indeed showed improved inhibition (62% vs 49%). When **I9G** was further modified with addition of a 2′-FRNA sugar at the -2 position forming **I12G**, we obtained our most potent A3G inhibitor with a *K*i of 670 ± 210 nM; the first nanomolar inhibitor of A3G-CTD2 to date. Interestingly, the hydrogen bond between D316 backbone NH with -1dC N3 (hydrogen bond 2, **Fig 4A**) appears to be strongly predictive of experimental inhibition and has previously been shown to be a critical A3G-ssDNA interaction(57). Overall, although our approach had more predictive value for A3G than for A3A, we show that pMD can be used to computationally guide the incorporation of sugar modifications into A3 targeting oligonucleotide inhibitors.

### Hairpin inhibitors with optimized PS patterns show single-digit nM potency against A3A

Prior characterization of A3A by us and other laboratories, including substrate-bound crystal structure (23,26,27), indicated that A3A prefers recognizing and binding to hairpin-shaped oligonucleotides. To further enhance binding, potency, and A3A targeting, we designed and evaluated a series of hairpin oligonucleotides utilizing a 13-nt self-complementary hairpin sequence with varying chemical modification patterns (**Fig 5**). As hairpin-structured, full PS oligonucleotide inhibitors have recently been shown to have equal potency to the equivalent PO inhibitor against A3A(30), we designed a series of four hairpins (**Fig 5A, B**) reflecting PS modification pattern findings from our linear inhibitors. Hairpin inhibitor **H1**_**A**_ is full PO DNA, and **H2**_**A**_ is its full PS equivalent (same backbone patterns as **I1**_**A**_ and **I2**_**A**_ respectively). **H3**_**A**_ is also full DNA, but contains PS linkages in all positions except at the linkages directly flanking the dZ (as for **I3**_**A**_), and **H4**_**A**_ is similar to **H3**_**A**_ but contains a terminal LNA sugar on both 5′ and 3′ ends. We hypothesized that the terminal LNA basepair in **H4**_**A**_ would provide additional hairpin stabilization and nuclease resistance relative to **H3**_**A**_.

**Figure 5.**
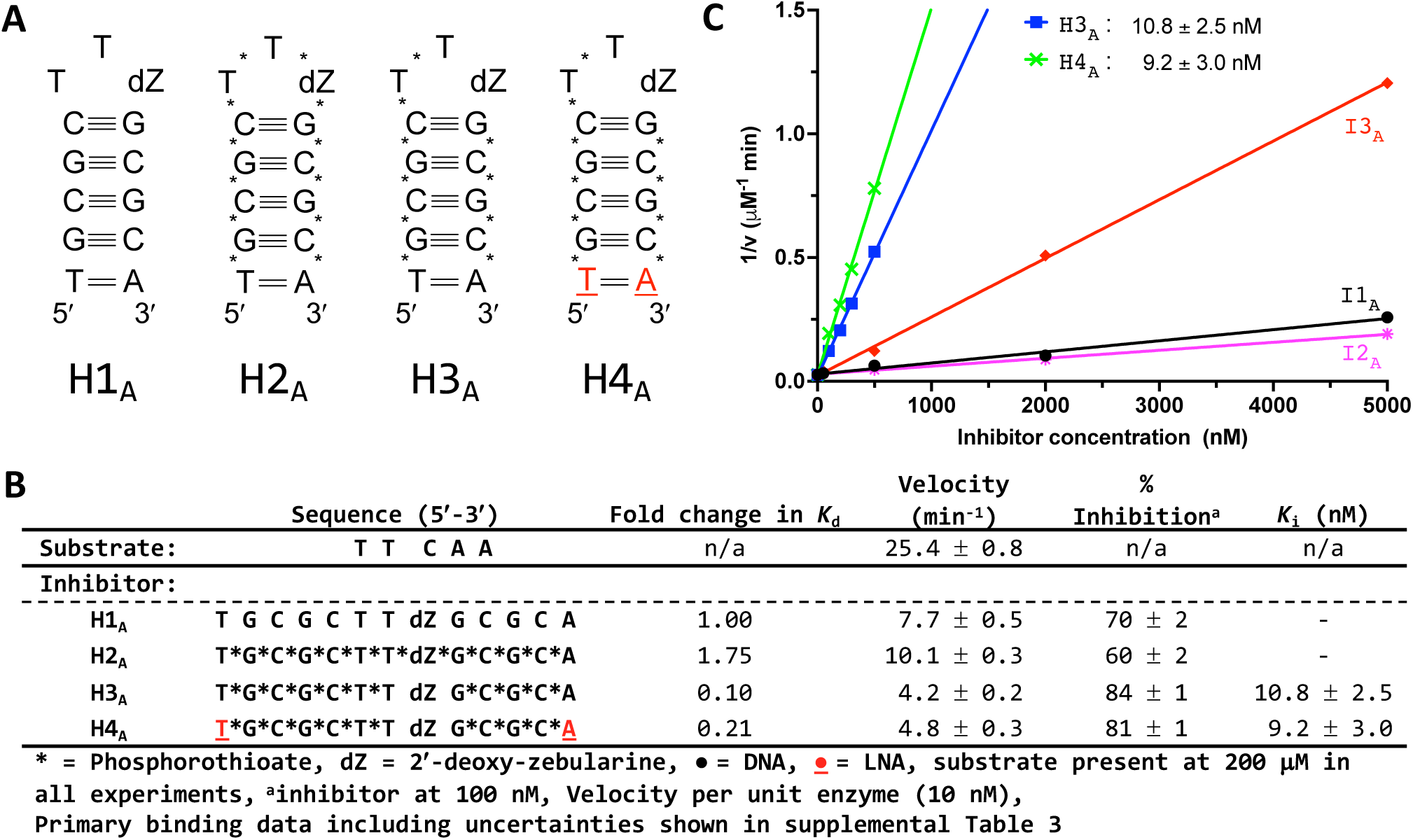
Hairpins with optimized chemical modifications are single-digit nM A3A inhibitors **A)** Hairpin sequence, structure, and chemical modification patterns targeting A3A (* denotes PS linkage, black is DNA sugar, and red is LNA sugar). **B)** Table of sugar and phosphate modified hairpin inhibitors targeting A3A. Relative binding affinities to A3A measured by MST, data shown as fold change relative to full PO control **H1** . Kinetic parameters vs A3A measured by ^1^H NMR deamination assay, percentage inhibition is relative to substrate only velocity. **C)** Inverse velocity vs inhibitor plot (Dixon plot) used to determine inhibition constants (*K*_i_) from the slope (see SI). Steeper line indicates a stronger inhibitor (datapoints triplicate, error bars SD).

We measured binding affinity using MST, and found that the hairpins demonstrated greatly enhanced A3A binding affinity relative to linear inhibitors, with the *K*d of full PO inhibitor **H1**_**A**_ being 5.2 ± 1.8 nM and full PS inhibitor **H2**_**A**_ being 9.1 ± 4.2 nM. Consistent with the results of our linear A3A substrates and inhibitors, removing the PS linkages flanking the active site increased binding affinity by an order of magnitude, giving a *K*d of 0.5 ± 0.1 nM and 1.1 ± 0.3 nM for **H3**_**A**_ and **H4**_**A**_ respectively. Next, we screened these four hairpins for inhibition activity in the ^1^H based NMR deamination assay at 100 nM concentration and confirmed **H3**_**A**_ and **H4**_**A**_ to have increased inhibition vs **H1**_**A**_ (84% and 81% vs 70%) (**Fig 5B**). Finally, we measured the *K*i for **H3**_**A**_ and **H4**_**A**_ and determined them to be highly potent, low nanomolar inhibitors of A3A (10.8 ± 2.5 nM and 9.2 ± 3.0 nM respectively, **Fig 5C**). These represent a ∼35-fold increase in potency compared to the linear unmodified **I1**_**A**_, and are the most potent inhibitors designed against A3A to date. Therefore, hairpin inhibitors containing optimized sugar- and phosphate-modification patterns resulted in significantly increased potency against A3A.

### Nuclease resistance of A3A inhibitors is proportional to PS content and structure

We evaluated the relative nuclease stability of our lead PS modified A3A inhibitors and compared them to the unmodified, full PO control inhibitors (**I1**_**A**_ and **H1**_**A**_). We incubated oligonucleotides in serum supplemented with fetal bovine serum (FBS), a source of 5′and 3′-exonucleases and endonucleases, as performed previously (31,58,59). The amount of remaining inhibitor over time was quantified by HPLC (**Fig 6)**.

**Figure 6.**
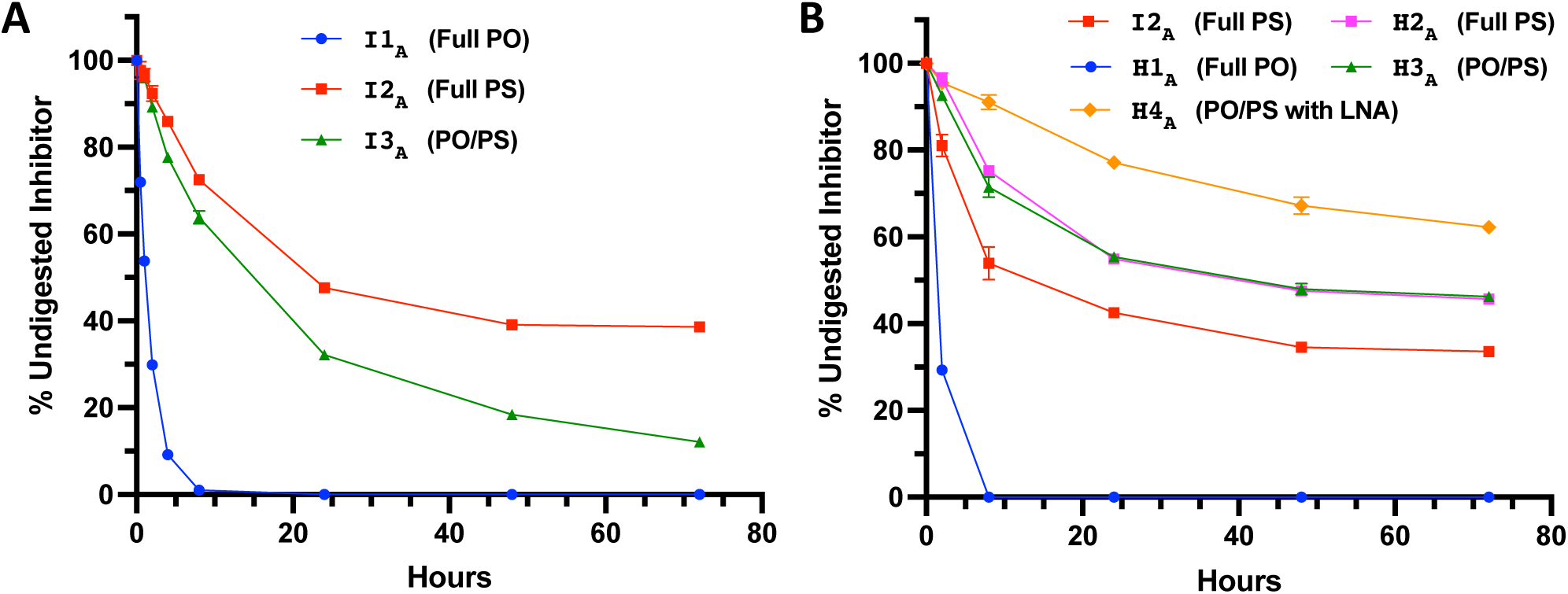
Nuclease stability of lead A3A inhibitors **A)** Serum stability data for linear A3A inhibitors **I1**_**A**_, **I2**_**A**_, and **I3**_**A**_ (datapoints duplicate, error bars SD). Serum stability data for hairpin-structured A3A inhibitors **H1**_**A**_, **H2**_**A**_, **H3**_**A**_, and **H4**_**A**_ with **I2**_**A**_ for comparison (datapoints duplicate, error bars SD).

For the linear inhibitors (**Fig 6A**), unmodified full PO **I1**_**A**_ was digested very quickly, with less than 10% present after 4 hours and was essentially undetectable after only 8 hours. In contrast, fully PS modified **I2**_**A**_, and partially PS modified **I3**_**A**_ (PO linkages flanking the target dZ only), were both dramatically more stable with 40% and 12% remaining respectively after 3 days. The lower stability of **I3**_**A**_ compared to **I2**_**A**_ is presumably due to **I3**_**A**_’s endonuclease susceptibility at the central unmodified PO linkages. While the fully PS-modified inhibitor **I2**_**A**_ was the most stable, the partially PS-modified inhibitor **I3**_**A**_ still retained significantly more stability than the unmodified inhibitor **I1**_**A**_, showing that unsurprisingly oligonucleotide nuclease stability is proportional to PS content.

The hairpin inhibitors were also evaluated using the same assay conditions (**Fig 6B**). The unmodified, full PO inhibitor **H1**_**A**_, was quickly digested within 8 hours. By contrast fully PS-modified **H2**_**A**_, and the partially modified **H3**_**A**_, plateaued at ∼50% undigested inhibitor after 3 days. Interestingly, the stability difference of **I2**_**A**_ vs **I3**_**A**_ is not seen for **H2**_**A**_ vs **H3**_**A**_ despite the same backbone PS pattern, suggesting the hairpin structure is likely protecting the central unmodified PO linkages within **H3**_**A**_ from endonuclease cleavage. Inhibitor **H4**_**A**_ showed the highest stability with 62% undigested inhibitor remaining after 72 hours validating that 5′ and 3′ terminal capping LNA sugars on **H4**_**A**_ provide significant additional hairpin stabilization and nuclease resistance. Thus, combining an optimal PS- and sugar-modification pattern into a hairpin structure, creates both the most potent A3A inhibitor, but also the most nuclease stable with the majority of inhibitor still present after 3 days incubating with serum.

## DISCUSSION

A3A and A3G are critical enzyme targets that have so far been essentially undruggable, and which contribute to tumor and viral sequence diversity that often leads to therapeutic escape. However, despite being a critical target and having a well-structured active-site, inhibition of A3 enzymes has been elusive, as attempts to develop small molecule inhibitors have been difficult. More recently, oligonucleotide-based inhibitors have been designed and tested against A3 enzymes and show promise, but still lack drug-like potency and cellular stability. In this study, we design chemically modified oligonucleotide inhibitors targeting A3A and A3G, leveraging a structure-based approach to direct the sites of chemical modification within the oligonucleotides.

Specifically, we interrogated the tolerance and impact of PS linkages on inhibition within these sequences for both A3 enzymes, and further investigated sugar modifications as a potential means to enhance inhibitor potency and stability. We discovered that for both A3 enzymes, due to the structural constraints of their active sites, inhibitors lost potency when PS modifications was incorporated at the two linkages directly flanking 5′ and 3′ the target dZ. However, careful positioning of PS linkages elsewhere in either sequence actually enhanced inhibitor potency. Mixed PO/PS backbones were favored for both enzymes proving to be the best linear inhibitors (*K*i’s: 1.43 μM against A3G (**I4G**) and 51 nM against A3A (**I3**_**A**_)), with natural achiral PO linkages flanking the target dZ and PS modifications elsewhere.

Sugar modifications incorporated into inhibitors targeting the less catalytically active and weaker binding A3G, also enhanced potency with *K*i of 670 nM (**I13G**), but had negligible impact on A3A inhibition. For A3A, we tested whether incorporating the mixed PO/PS backbone enhanced binding affinity and inhibition further in the context of its preferred structural substrate, a DNA hairpin. Use of a fully PS-modified hairpin inhibitor was recently shown by Harjes *et al.*(30) to provide additional nuclease stability, but also showed a slight loss of inhibition relative to a full PO inhibitor, similar to our results for both A3 enzymes. In contrast, we discovered that incorporation of our optimal PO/PS backbone pattern into the A3A targeting hairpin sequence resulted in our most potent inhibitors We also evaluated nuclease stability of our A3A-targeting inhibitors by incubation with serum supplemented media. Not surprisingly, fully PO inhibitors were rapidly and completely degraded within 4-8 hours, however PS modifications offered drastically higher nuclease stability on the order of days. PS modified linear inhibitors (**I2**_**A**_ and **I3**_**A**_) remained ∼40% and ∼12% undigested at 3 days, and this was further improved to around ∼50% for hairpin structured inhibitors (**H2**_**A**_ and **H3**_**A**_). Excitingly, our lead A3A inhibitor containing optimized PO/PS backbone modification pattern with further stabilization of the hairpin using 5′ and 3′ terminal LNA sugar modifications resulted in the most nuclease resistant (>60% undigested after 3 days), and most potent A3A inhibitor **H4**_**A**_ (*K*i of 9.2 nM) reported to date. Our lead A3A inhibitors were further validated in a cellular A3A inhibition study. While the inhibitors with exclusively PO had no inhibitory effect, the linear PS showed some inhibition, and the chemically modified hairpins showed significant inhibition of A3A in a cellular environment. Thus, we have demonstrated that leveraging knowledge of both the structural biology of A3-complexes, and chemical modification of oligonucleotides, have led to compounds with remarkable potency, stability, and activity within cells. Based on these findings, our expectation is that we can design cellularly stabilized A3 specific inhibitors, based on the substrate specificity of the particular A3 enzyme.

While our study shows that nucleic acid modifications are transformative for both potency and cellular stability of A3 targeting oligonucleotide inhibitors, this work has focused only on a subset of common nucleic acid modifications. Further optimization of sugarand phosphatemodifications and transitionstate mimics in A3 targeting oligonucleotide inhibitors are likely to lead to further increases in enzyme specificity, inhibitor potency, and nuclease stability. Most likely, chemical modification of these inhibitors in positions flanking the target dZ requires additional focus. For example, we used traditional diastereomeric PS linkages which are a ∼1:1 mix of *R*p and *S*p-centers at phosphorus, however despite its increased synthetic complexity, chirally-controlled PS synthesis is gaining traction(60–62). When targeting proteins with PS-modified oligonucleotides, it has been demonstrated previously from structural studies there are preferences for particular PS diastereomers at specific positions (63–65). With this in mind we modeled 4 diastereomeric PS combinations (i.e. *RR*, *RS*, *SR*, and *SS*) in the positions flanking the active site, however these preliminary models appeared to be unsuccessful **(Supplementary Fig S4).** Nevertheless, modulation of PS chirality in A3 oligonucleotide inhibitors may be able to confer additional binding and specificity.

Development of cellularly stable and potent A3 specific inhibitors is noteworthy as the ultimate therapeutic potential of bioavailable A3 specific inhibitors is significant. For many human cancers and viral infections A3 enzymes are the source of heterogeneity, and each A3 enzyme has a unique sequence signature, or structure that they recognize, as they catalyze the mutation of C to U in the host or viral genome(8). In cancer therapy, mutational heterogeneity leads to drug resistance and therapeutic tolerance(66). Mutations introduced from the most catalytically active A3A, which recognizes a tight hairpin structure contributes to the largest numbers of mutations in a wide variety of cancers, while A3B also contributes and recognizes a more extended hairpin(67) has now also been observed to be a mutational driver(68). The less active A3G contributes to diversity in viral infections in HIV(6,69,70) and HTLV(71) and has also been recently implicated in the clonal diversity of bladder cancer(72), multiple myeloma(73) as well as effects in other cancer genomes(74,75). With drug resistance preventing the realization of the therapeutic potential of many chemotherapeutic and antiviral drugs, co-inhibiting A3 enzymes is likely a viable strategy to restrict the speed by which mutations confer drug resistance in both oncology and infectious disease.

## SUPPLEMENTARY DATA

PDF available online containing dZ phosphoramidite synthesis details, scheme, and NMRs; oligonucleotide characterization, protein expression details, and MST binding values.

## AUTHOR CONTRIBUTIONS

CAS and AKH conceived the study with help from HM and JKW. AKH performed dZ phosphoramidite synthesis; oligonucleotide design, synthesis, and purification; A3G-CTD2 NMR inhibition experiments; nuclease/HPLC experiments; and formal data analysis. WM and VB performed A3A and A3A-E72A expression and purification; and MST binding experiments. WM performed A3A NMR inhibition experiments. DS performed modelling; molecular dynamics simulations; and data analysis. AMS performed A3G-CTD2 expression and purification. NKY and JML helped with data analysis. CAS, HM, JKW, NKY and JML supervised the research. AKH and CAS wrote the manuscript with input and editing from all other authors.

## ACKNOWLEDGMENTS

The authors thank Dr. Jasna Fejzo and the UMass Amherst NMR Core facility for help and guidance with A3G NMR inhibition experiments. We thank Dr. Janusz Koscielniak for maintenance of the NMR spectrometers in the NMR Facility for Biological Research, Center for Structural Biology, Center for Cancer Research, National Cancer Institute, and the Biophysics Resource in the Center for Structural Biology, Center for Cancer Research, National Cancer Institute for assistance with MST measurements.

