## Supplementary Information for "Next generation APOBEC3 inhibitors: Optimally designed for potency and nuclease stability"

**General Chemistry:**

Chemicals, reagents, and anhydrous solvents were purchased from commercial sources (Sigma Aldrich, Chemgenes, Biosynth) and were used as is, unless otherwise stated. All reactions were performed in oven-dried round bottomed flasks fitted with rubber septa under dry Ar atmosphere, unless otherwise noted. Compounds were purified by flash chromatography (Biotage prepacked silica cartridges). Analytical thin-layer chromatography (TLC) was performed using silica gel (60 F-254) coated aluminum plates (Merck), and spots were visualized by exposure to ultraviolet light (UV), and/ or stained with 10 % $H_2SO_4$ in ethanol or ninhydrin, followed by brief heating. $^{1}$H, $^{13}$C{1H},$^{135}$DEPT, $^{31}$P, and 2D NMR spectra were acquired on a Bruker Avance IIIHD 500 MHz NMR instrument. Chemical shifts are reported in ppm (*δ* scale) relative to the solvent signal and coupling constant (*J*) values are reported in hertz. Data are presented as follows: chemical shift, multiplicity; ((s) singlet, (d) doublet, (t) triplet, (q) quartet, (qu) quintet, (m) multiplet, (br) broad, (dd) doublet of doublets, (dt) doublet of triplets), coupling constant in Hz, and integration. Structural assignments were made with additional information from gCOSY, gHSQC, gHMBC, and gNOESY experiments as needed. For High-Resolution Mass Spectrometry, samples were re-suspended in acetonitrile, methanol, or water, and analyzed by flow-injection analysis using a Thermo Scientific Orbitrap Velos Pro mass spectrometer coupled to a Thermo Accela 1250 UPLC.

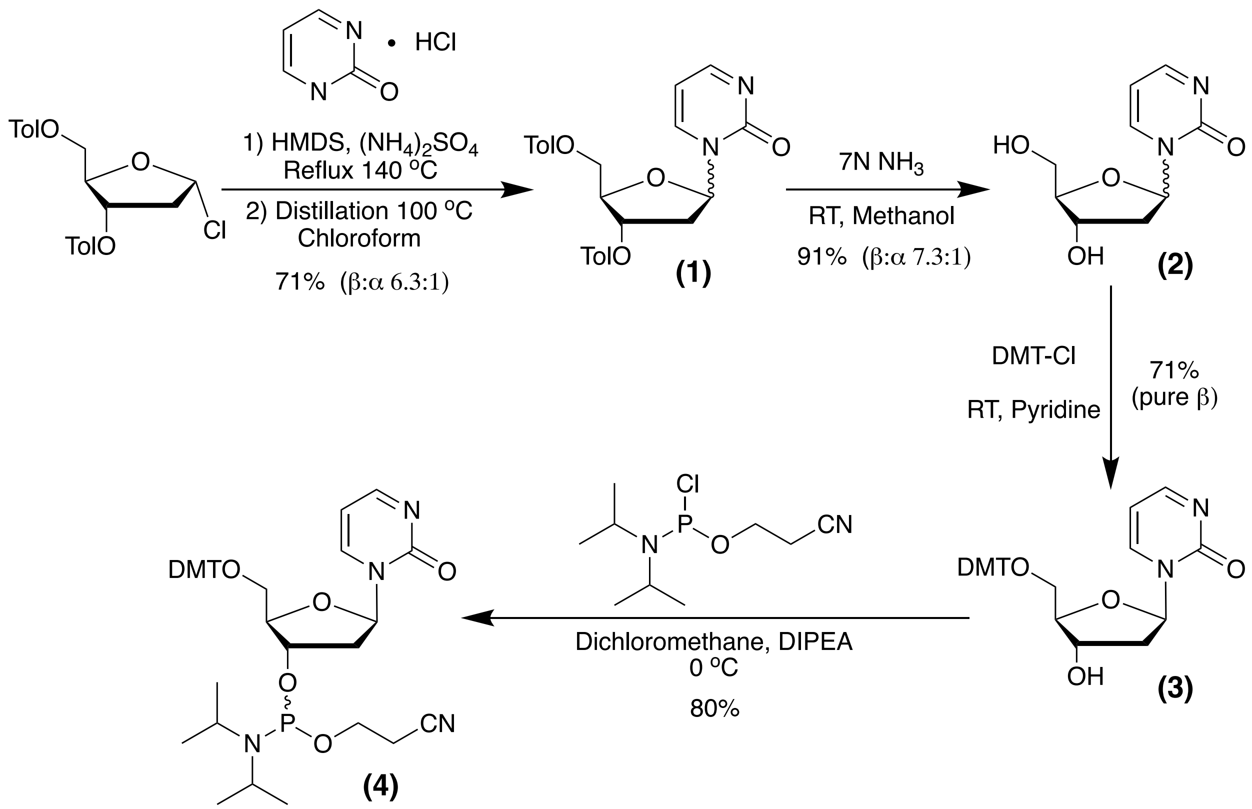

**Scheme S1: Synthetic route to dZ-phosphoramidite**

**dZ phosphroramidite yield and comparison to Harjes *et al.* (*Biochemistry 2019, 58, 5, 391–400*)**

- Total phosphoramidite synthesis yield herein (71% x 91% x 71% x 80%) = **<u>37%</u>**
- Total phosphoramidite synthesis yield reported previously Harjes *et al.* (91% x 54% x 54% x 88%) = **<u>24%</u>**

- Improvements compared to Harjes *et al.* procedure; removing fractional distillation of the silylated nucleobase in favor of simple evaporation, and no need for column purification of (1) post glycosylation – thorough extraction is sufficient. Removal of α-anomer of (1) based on its insolubility in 40% hexane in chloroform at -20 °C overnight, gives white cotton-wool like precipitate, to enrich the anomeric ratio.

*This represents a ~50% increase in yield with added improvements to synthetic simplicity*

**NMR tabulation for compounds (1) to (4):**

**(1 - β anomer)** - 3′-5′-Di-O-(p-toluoyl)-2'-deoxyzebularine (6.3:1 β:α, after α-anomer removal):

**$^{1}$H NMR** (500 MHz, DMSO-$d_6$): $\delta$ 8.57 (dd, $J$ = 4.1, 2.8 Hz, 1H, Ar-CH), 8.27 (dd, $J$ = 6.8, 2.8 Hz, 1H, Ar-CH), 7.94-7.92 (m, 2H, Ar-CH), 7.81-7.78 (m, 2H, Ar-CH), 7.36 (d, $J$ = 8.1 Hz, 2H, Ar-CH), 7.30 (d, $J$ = 8.1 Hz, 2H, Ar-CH), 6.45 (dd, $J$ = 6.7, 4.1 Hz, 1H, Ar-CH), 6.22 (t, $J$ = 6.7 Hz, 1H, CH), 5.59 (m, 1H, CH), 4.69-6.66 (m, 1H, CH), 4.64-6.61 (m, 2H, $CH_2$), 2.85-80 (ddd, $J$ = 14.6, 6.3, 2.7 Hz, 1H, $CH_2$), 2.53-4.49 (m, 1H, $CH_2$ (overlapping DMSO)), 2.40 (s, 3H, Ar-$CH_3$), 2.37 (s, 3H, Ar-$CH_3$)

**$^{13}$C NMR** (126 MHz, DMSO-$d_6$): $\delta$ 166.42, 165.45, 165.23, 154.56, 144.09, 143.93, 129.48, 129.34, 129.33, 129.23, 126.46, 126.43, 104.07, 87.82, 82.65, 79.17, 74.84, 64.21, 37.92, 21.21, 21.16

**(1 - α -anomer)** - 3′-5′-Di-*O*-(p-toluoyl)-2'-deoxyzebularine (α-anomer):

**$^{1}$H NMR** (500 MHz, DMSO-$d_6$): $\delta$ 8.58 (dd, $J$ = 4.1, 2.8 Hz, 1H, Ar-CH), 8.43 (dd, $J$ = 6.7, 2.8 Hz, 1H, Ar-CH), 7.95-7.92 (m, 2H, Ar-CH), 7.61-7.58 (m, 2H, Ar-CH), 7.38 (d, $J$ = 8.1 Hz, 2H, Ar-CH), 7.27 (d, $J$ = 8.1 Hz, 2H, Ar-CH), 6.47 (dd, $J$ = 6.7, 4.1 Hz, 1H, Ar-CH), 6.15 (d, $J$ = 6.4 Hz, 1H, CH), 5.54 (d, $J$ = 5.9 Hz, 1H, CH), 5.17 (t, $J$ = 4.9 Hz, 1H, CH), 4.53-4.47 (m, 2H, $CH_2$), 2.97 (dt, $J$ = 15.4, 6.4 Hz, 1H, $CH_2$), 2.40 (s, 3H, Ar-$CH_3$), 2.38 (s, 3H, Ar-$CH_3$)

**(2 - β anomer)** - 2′-deoxyzebularine (6.3:1 β:α)

**$^{1}$H NMR** (500 MHz, DMSO-$d_6$): $\delta$ 8.56 (dd, $J$ = 4.1, 2.9 Hz, 1H, Ar-CH), 8.47 (dd, $J$ = 6.7, 2.8 Hz, 1H, Ar-CH), 6.50 (dd, $J$ = 6.7, 4.2 Hz, 1H, Ar-CH), 6.07 (d, $J$ = 6.2 Hz, 1H, CH), 5.30 (br s, 1H, OH), 5.10 (br s, 1H, OH), 4.24-4.20 (m, 1H, CH), 3.90 (q, $J$ = 3.7 Hz, 1H, CH), 3.62 (ddd, $J$ = 20.1, 15.5, 3.5 Hz, 2H, $CH_2$), 2.40-2.34 (m, 1H, $CH_2$), 2.05-2.00 (m, 1H, $CH_2$)

**$^{13}$C NMR** (126 MHz, DMSO-$d_6$): $\delta$ 165.87, 154.68, 144.15, 103.79, 88.12, 86.77, 69.63, 61.67, 60.68, 41.03

**(3)** - 5′-*O*-(4,4′-dimethoxytrityl)-2′-deoxyzebularine (Pure β anomer)

**$^{1}$H NMR** (500 MHz, DMSO-$d_6$): $\delta$ 8.55 (dd, $J$ = 4.1, 2.8 Hz, 1H, Ar-CH), 8.22 (dd, $J$ = 6.7, 2.8 Hz, 1H, Ar-CH), 7.39-7.29 (m, 4H, Ar-CH), 7.28-7.21 (m, 5H, Ar-CH), 6.93-6.88 (m, 4H, Ar-CH), 6.25 (dd, $J$ = 6.7, 4.1 Hz, 1H, Ar-CH), 6.07 (d, $J$ = 5.9 Hz, 1H, CH), 5.39 (d, 1H, OH), 4.29 (qu, $J$ = 5.3 Hz, 1H, CH), 4.01 (q, $J$ = 4.1 Hz, 1H, CH), 3.74 (s, 6H, Ar-$CH_3$), 3.31-3.25 (m, 2H, $CH_2$), 2.45-2.39 (m, 1H, $CH_2$), 2.16-2.10 (m, 1H, $CH_2$)

$^{13}$**C NMR** (126 MHz, DMSO-$d_6$): $\delta$ 165.98, 158.11, 154.52, 144.57, 143.65, 135.31, 135.12, 129.72, 129.70, 127.89, 127.65, 126.77, 113.24, 103.61, 86.70, 85.98, 85.88, 79.15, 69.31, 62.76, 55.03, 40.83

**(4)** - 5′-*O*-(4,4′-dimethoxytrityl)-2′-deoxyzebularine,3'-[(2-cyanoethyl)-(*N*,*N*-diisopropyl)]-phosphoramidite (Pure β anomer, ~1:1 diastereomers)

The mixture of two (~1:1) diastereomers at phosphorus led to complex $^{1}$H and $^{13}$C NMR spectra, full spectra are shown below.

$^{31}$**P NMR** (202 MHz, DMSO-$d_6$): $\delta$ 147.81 (sextet, $J$ = 9.0 Hz, 1P), 147.52 (sextet, $J$ = 9.0 Hz, 1P)

$^{31}$**P$^{\{1H\}}$ NMR** (202 MHz, DMSO-$d_6$): $\delta$ 147.81, 147.52 and small P(v) impurity (H-phosphonate) at 13.86

**NMR spectra for compounds (1) to (4):**

**(1)** - 3′-5′-Di-*O*-(p-toluoyl)-2'-deoxyzebularine (6.3:1 β:α, after α-anomer removal, and without column purification):

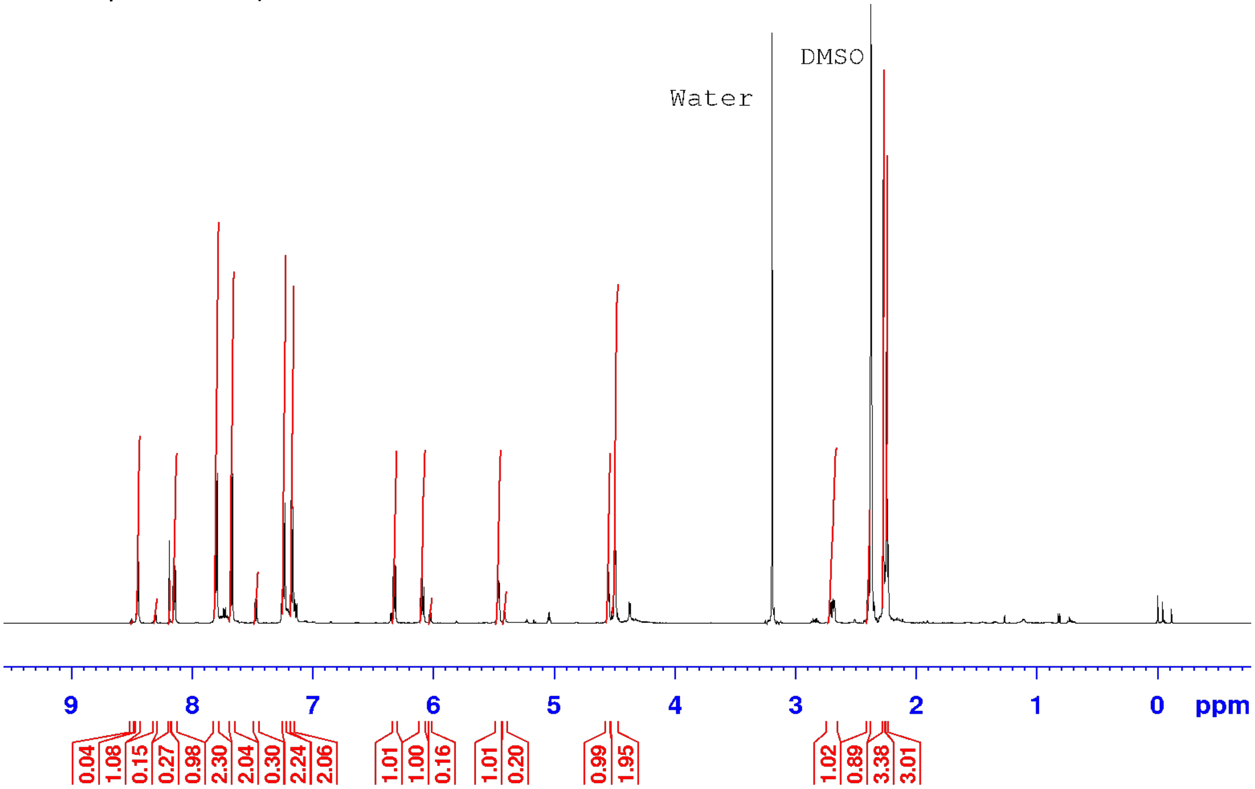

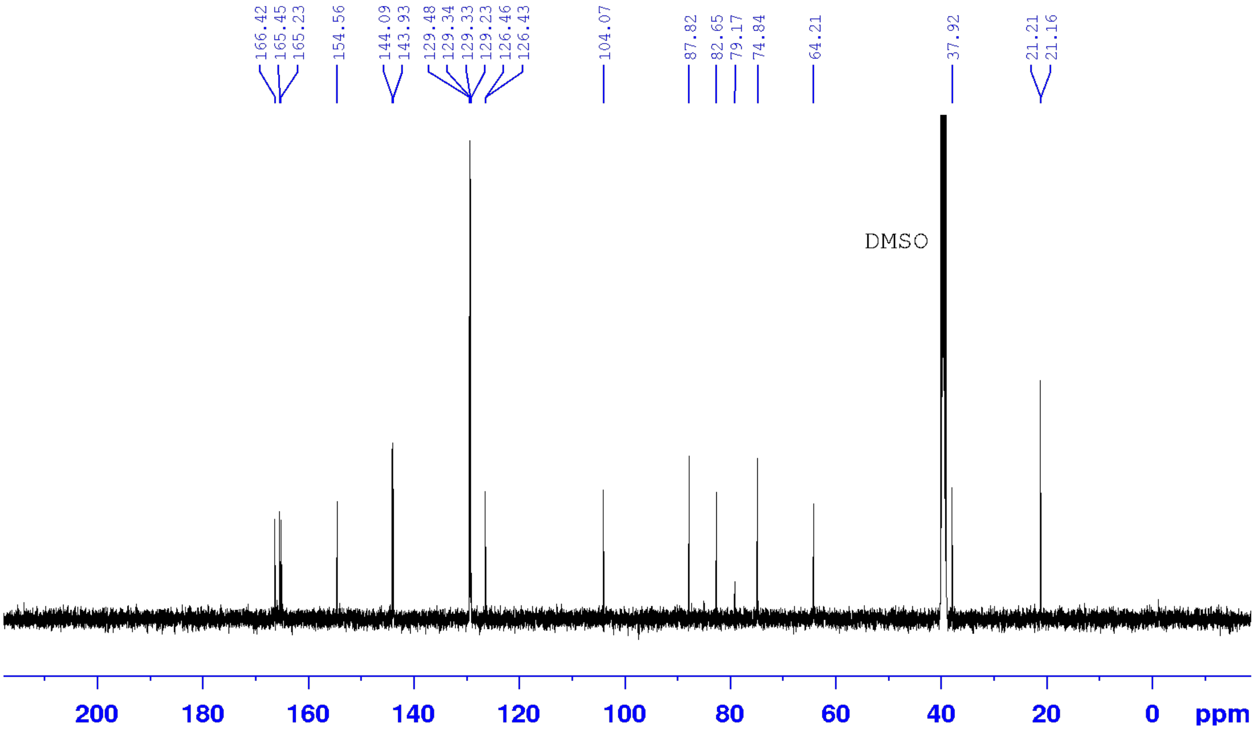

**(1)** - 3′-5′-Di-*O*-(p-toluoyl)-2'-deoxyzebularine ($\alpha$-anomer):

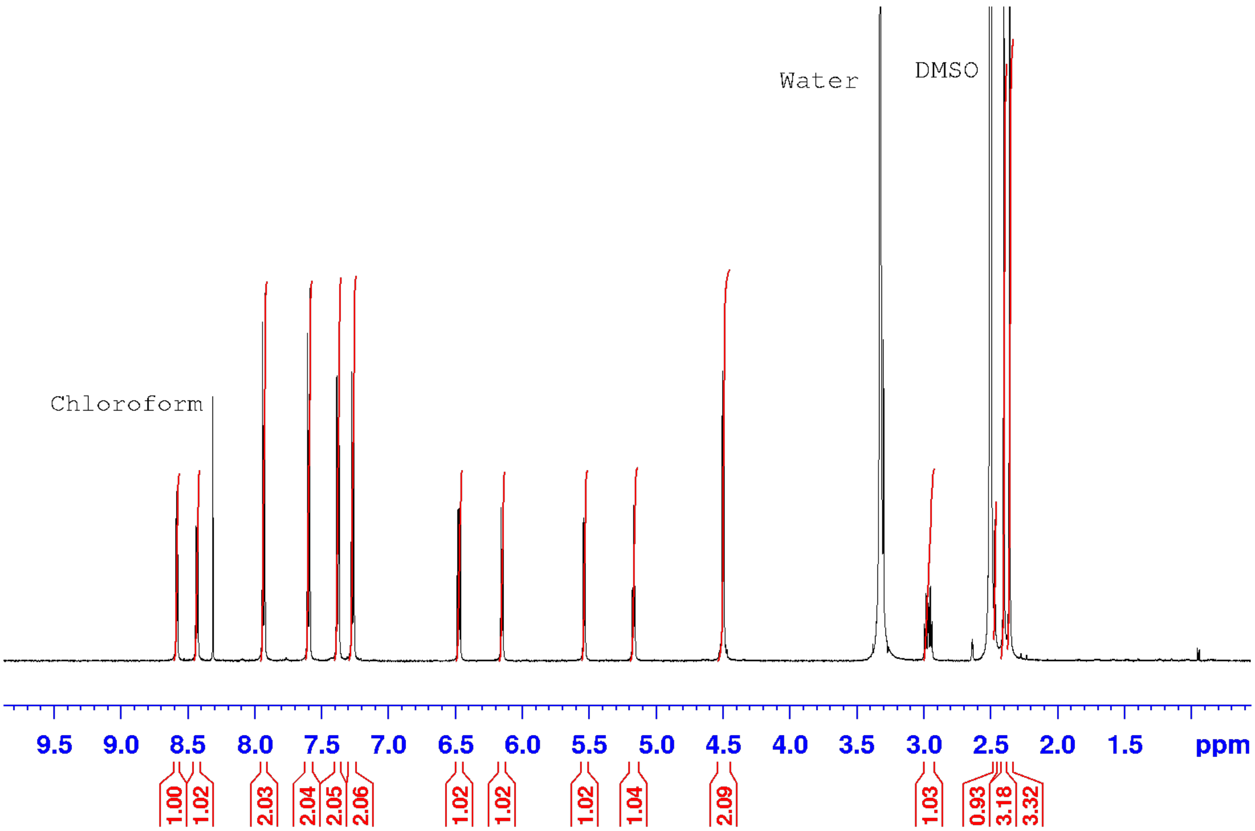

**(2)** - 2′-deoxyzebularine (6.3:1 β:α, without column purification):

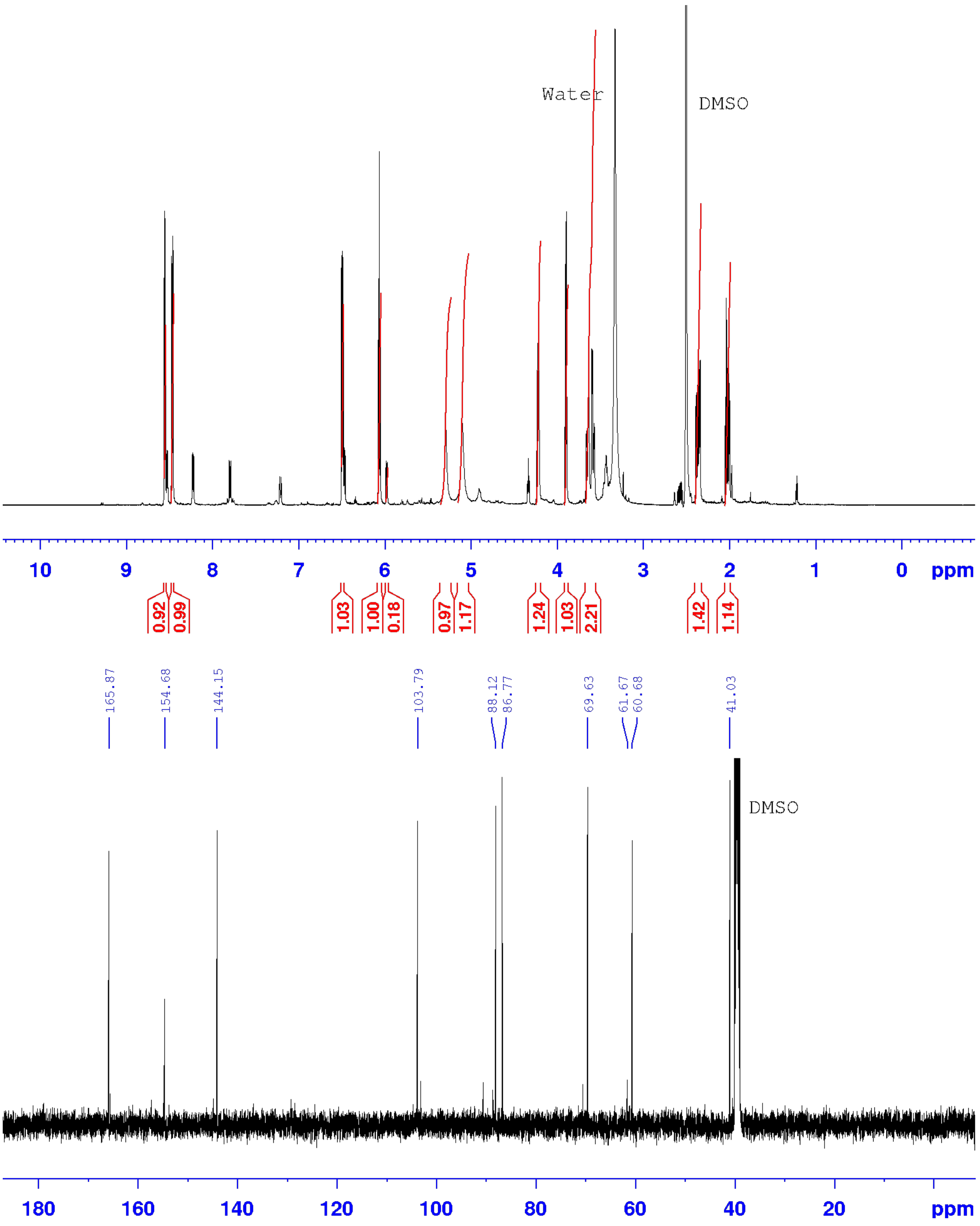

**(3)** - 5′-*O*-(4,4′-dimethoxytrityl)-2′-deoxyzebularine (Pure β anomer):

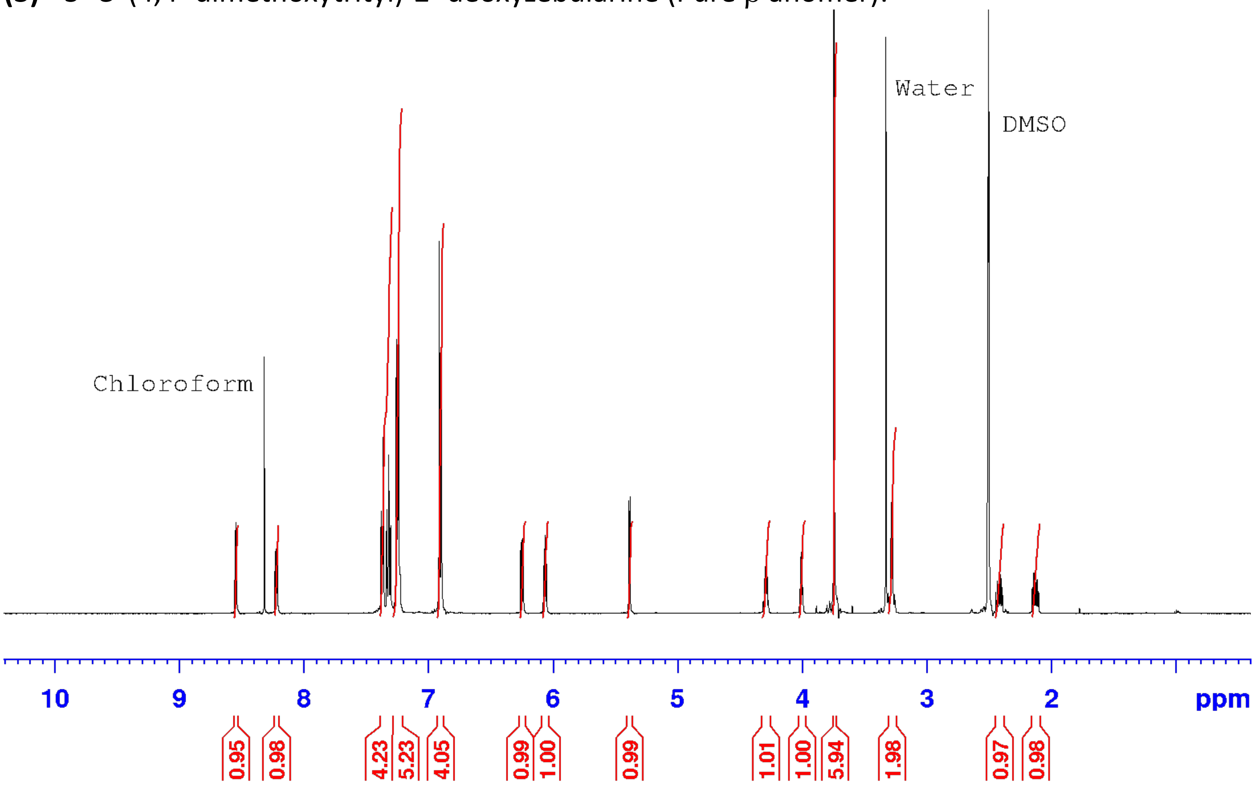

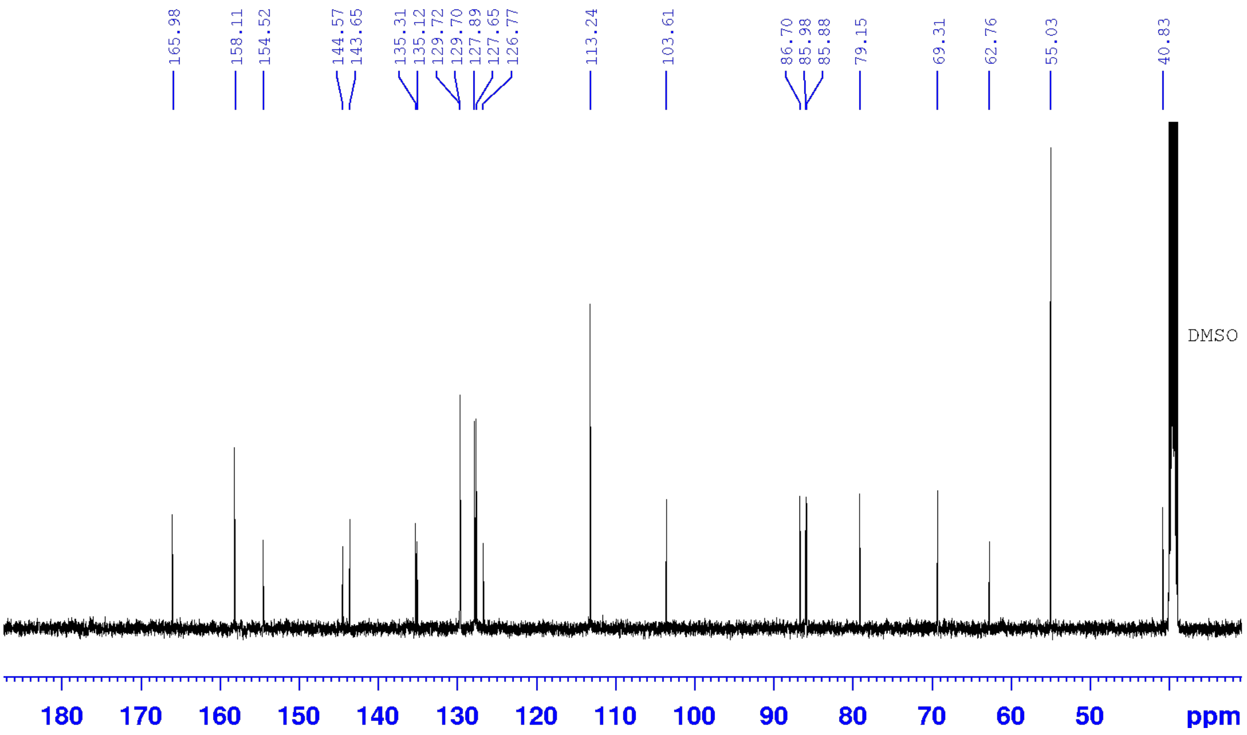

**(4)** - 5′-*O*-(4,4′-dimethoxytrityl)-2′-deoxyzebularine,3'-[(2-cyanoethyl)-(*N*,*N*-diisopropyl)]-phosphoramidite (Pure β anomer, ~1:1 diastereomers)

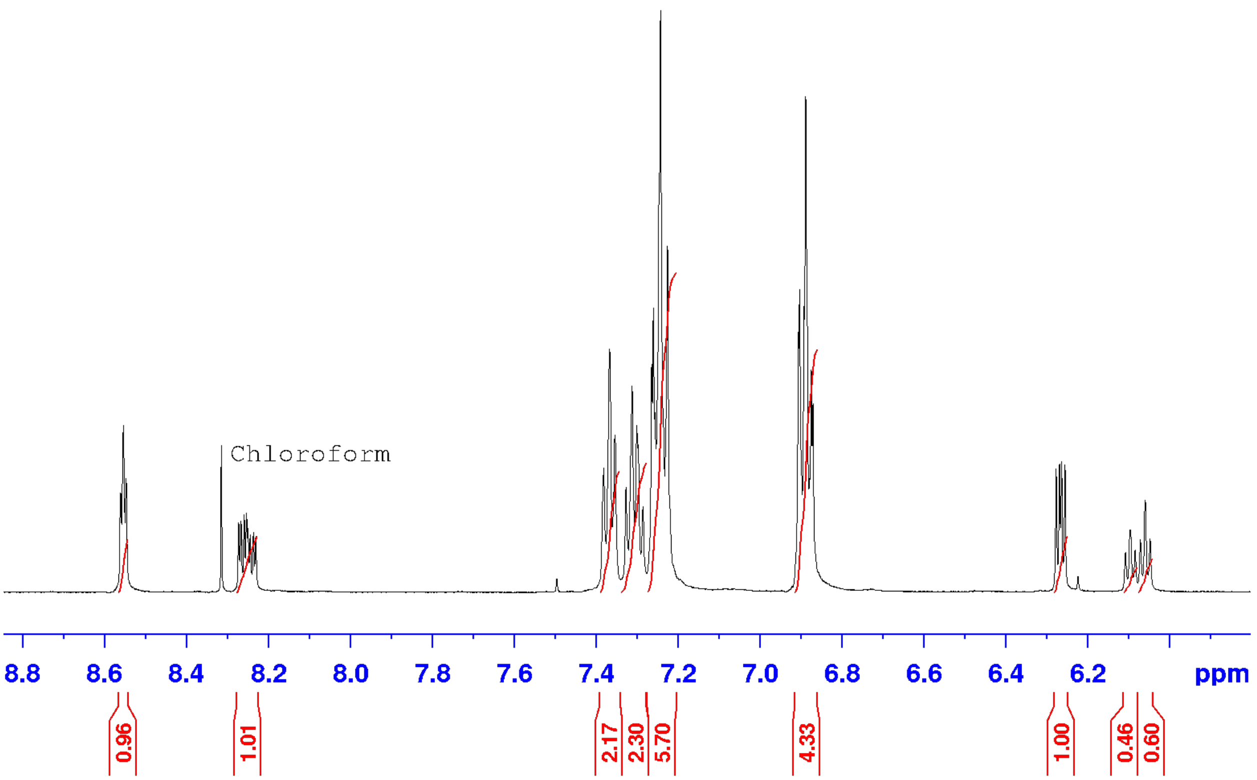

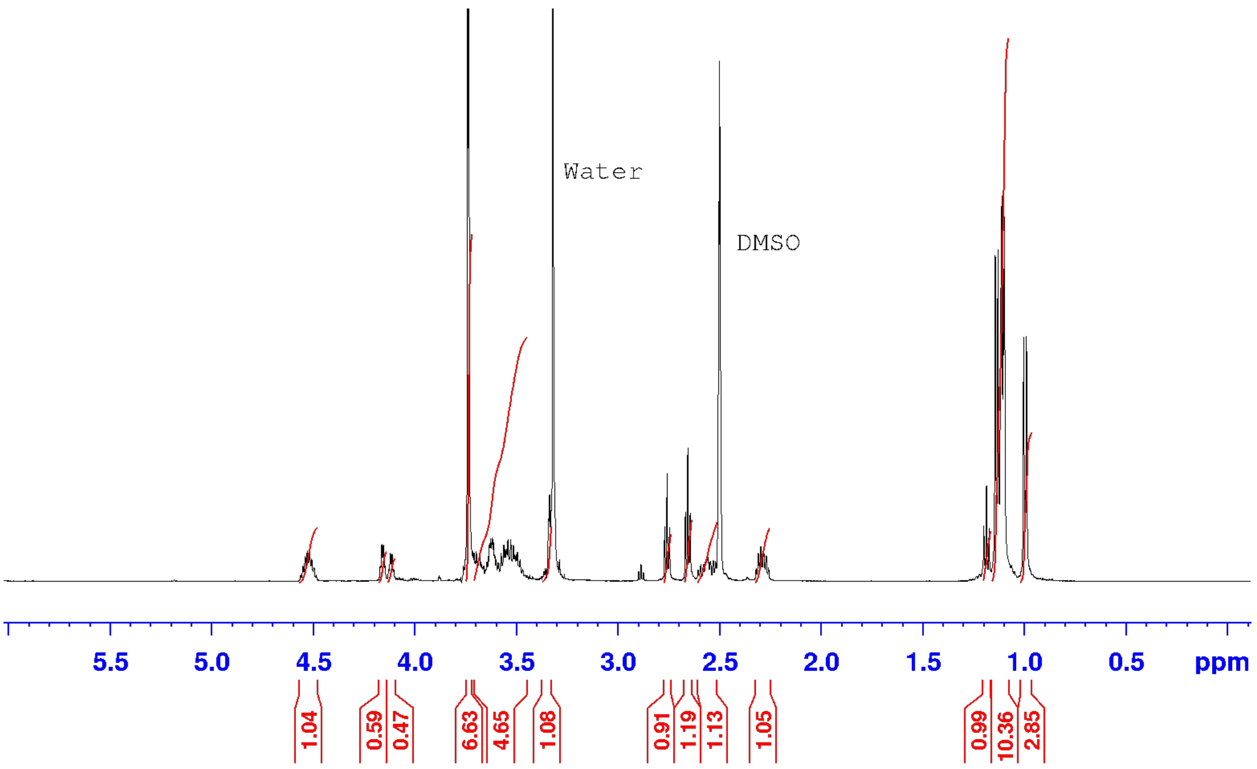

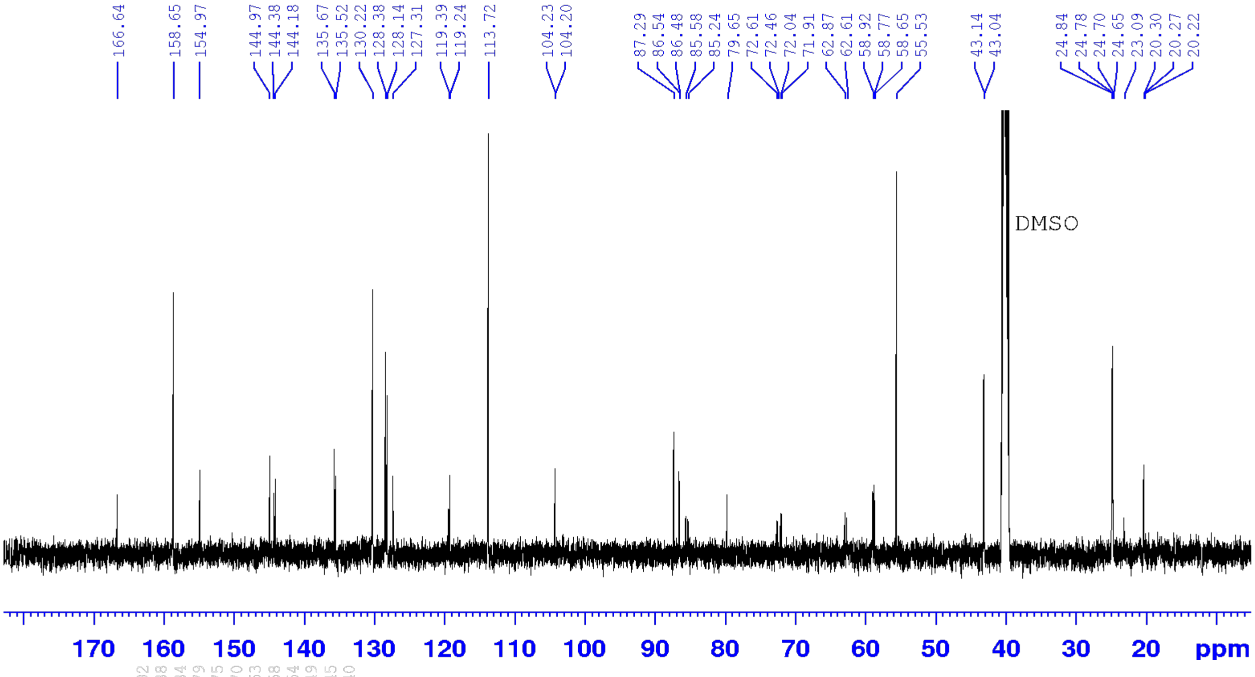
166.64
158.65
154.97
144.97
144.38
144.18
135.67
135.52
130.22
128.38
128.14
127.31
119.39
119.24
113.72
104.23
104.20
87.29
86.54
86.48
85.58
85.24
79.65
72.61
72.46
72.04
71.91
62.87
62.61
58.92
58.77
58.65
55.53
43.14
43.04
24.84
24.78
24.70
24.65
23.09
20.30
20.27
20.22
DMSO
170 160 150 140 130 120 110 100 90 80 70 60 50 40 30 20 ppm

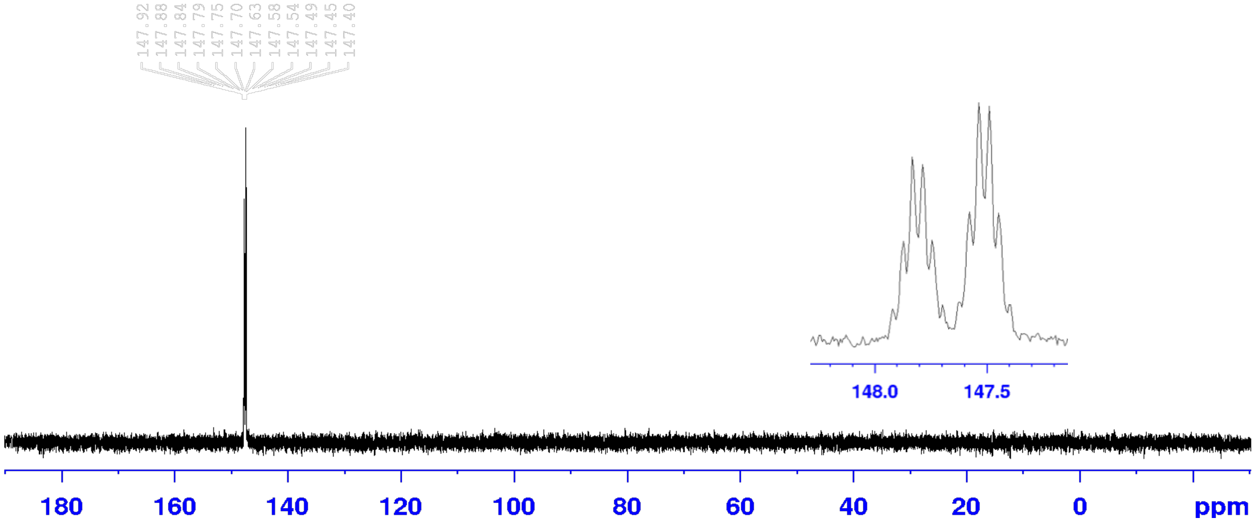
147.92
147.88
147.84
147.79
147.75
147.70
147.63
147.58
147.54
147.49
147.45
147.40
148.0
147.5
180 160 140 120 100 80 60 40 20 0 ppm

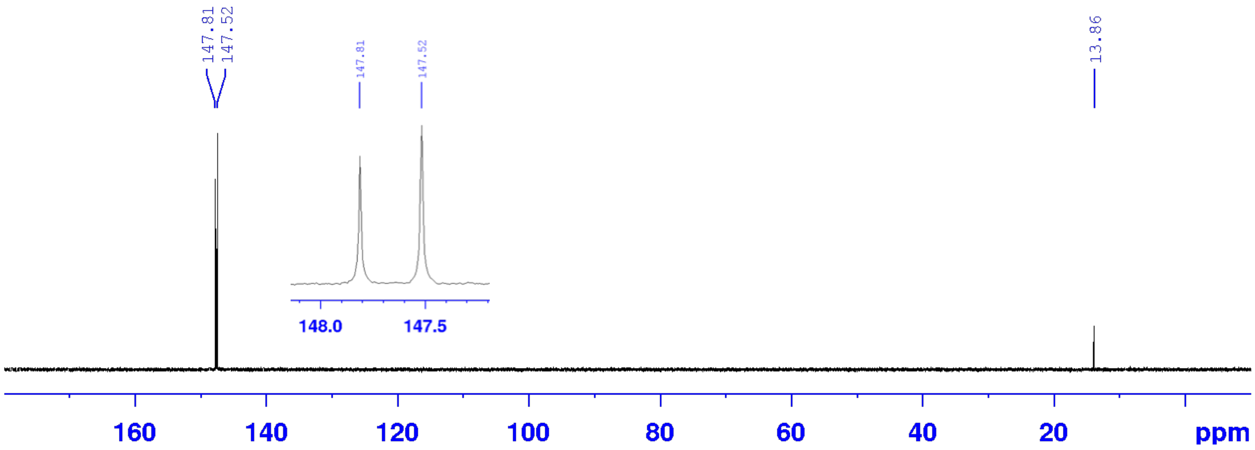
147.81
147.52
13.86
148.0
147.5
160 140 120 100 80 60 40 20 ppm

| Name | Sequence (5′-3′) | Calc. Epsilon | Calc. MW | Found MW |
|---|---|---|---|---|
| I1$_G$ | (dA)(dA)(dT)(dC)(dC)(dZ)(dA)(dA)(dA) | 90900 | 2660.79 | 2660.50 |
| I2$_G$ | (dA)*(dA)*(dT)*(dC)*(dC)*(dZ)*(dA)*(dA)*(dA) | 90900 | 2789.32 | 2788.30 |
| I3$_G$ | (dA)(dA)(dT)*(dC)*(dC)*(dZ)*(dA)*(dA)(dA) | 90900 | 2741.12 | 2740.40 |
| I4$_G$ | (dA)*(dA)*(dT)(dC)(dC)(dZ)(dA)*(dA)*(dA) | 90900 | 2725.06 | 2724.41 |
| I5$_G$ | (dA)*(dA)*(dT)*(dC)*(dC)(dZ)(dA)*(dA)*(dA) | 90900 | 2757.19 | 2756.37 |
| I6$_G$ | (dA)(dA)(dT)(dC)(dC)*(dZ)*(dA)(dA)(dA) | 90900 | 2692.93 | 2692.42 |
| I7$_G$ | (fA)(fA)(fT)(xC)(fC)(dZ)(fA)(fA)(dA) | 90720 | 2796.75 | 2796.43 |
| I8$_G$ | (dA)(dA)(fT)(xC)(fC)(dZ)(dA)(fA)(dA) | 90720 | 2742.77 | 2742.45 |
| I9$_G$ | (dA)(dA)(fT)(dC)(dC)(dZ)(fA)(dA)(dA) | 90900 | 2696.77 | 2696.48 |
| I10$_G$ | (dA)*(dA)*(fT)(dC)(dC)(dZ)(fA)*(dA)*(dA) | 90900 | 2761.04 | 2760.39 |
| I11$_G$ | (dA)(dA)(dT)(fC)(fC)(dZ)(dA)(dA)(dA) | 90540 | 2696.77 | 2696.45 |
| I12$_G$ | (dA)(dA)(dT)(wC)(dC)(dZ)(dA)(dA)(dA) | 90720 | 2678.78 | 2678.48 |
| I13$_G$ | (dA)(dA)(fT)(wC)(dC)(dZ)(fA)(dA)(dA) | 90720 | 2714.76 | 2714.46 |
| I1$_A$ | (dA)(dA)(dA)(dT)(dT)(dZ)(dA)(dA)(dA)(dA)(dA)(dA)(dA) | 154710 | 3952.64 | 3951.75 |
| I2$_A$ | (dA)*(dA)*(dA)*(dT)*(dT)*(dZ)*(dA)*(dA)*(dA)*(dA)*(dA)*(dA)*(dA) | 154710 | 4145.44 | 4144.45 |
| I3$_A$ | (dA)*(dA)*(dA)*(dT)*(dT)(dZ)(dA)*(dA)*(dA)*(dA)*(dA)*(dA)*(dA) | 154710 | 4113.31 | 4122.49 |
| I4$_A$ | (dA)(dA)(dA)(dT)(dT)*(dZ)*(dA)(dA)(dA)(dA)(dA)(dA)(dA) | 154710 | 3984.77 | 3984.69 |
| I5$_A$ | (dA)*(dA)*(dA)*(dT)(dT)(dZ)(dA)(dA)*(dA)*(dA)*(dA)*(dA)*(dA) | 154710 | 4081.17 | 4080.56 |
| I6$_A$ | (dA)(dA)(dA)(dT)*(dT)*(dZ)*(dA)*(dA)(dA)(dA)(dA)(dA)(dA) | 154710 | 4016.91 | 4015.65 |
| I7$_A$ | (dA)(dA)(dA)(dT)(fT)(dZ)(fA)(dA)(dA)(dA)(dA)(dA)(dA) | 154710 | 3988.62 | 3987.73 |
| I8$_A$ | (dA)(dA)(dA)(fT)(dT)(dZ)(dA)(dA)(dA)(dA)(dA)(dA)(dA) | 154710 | 3970.63 | 3970.74 |
| H1$_A$ | (dT)(dG)(dC)(dG)(dC)(dT)(dT)(dZ)(dG)(dC)(dG)(dC)(dA) | 105840 | 3911.55 | 3910.66 |
| H2$_A$ | (dT)*(dG)*(dC)*(dG)*(dC)*(dT)*(dT)*(dZ)*(dG)*(dC)*(dG)*(dC)*(dA) | 105840 | 4104.35 | 4103.37 |
| H3$_A$ | (dT)*(dG)*(dC)*(dG)*(dC)*(dT)*(dT)(dZ)(dG)*(dC)*(dG)*(dC)*(dA) | 105840 | 4072.22 | 4071.42 |
| H4$_A$ | (lT)*(dG)*(dC)*(dG)*(dC)*(dT)*(dT)(dZ)(dG)*(dC)*(dG)*(dC)*(lA) | 105840 | 4128.24 | 4127.41 |
| Key: * = phosphorothioate, dZ = deoxyzebularine, (dx) = DNA, (fx) = 2′-FANA, (wx) = 2′-FRNA, (lx) = LNA | | | | |

**Table S1: Oligonucleotide inhibitors synthesized in this work with calculated epsilon values, calculated molecular weights, and LCMS confirmed molecular weights**

| Oligo | Sequence (5′-3′) | Binding affinity $K_d$ (nM) Substrate (X = dC) | Inhibitor (X = dZ) |
|---|---|---|---|
| S1$_A$ or I1$_A$ | A A A T T X A A A A A A A | 20 ± 4 | 141 ± 87 |
| S2$_A$ or I2$_A$ | A*A*A*T*T*X*A*A*A*A*A*A*A | 25 ± 5 | 233 ± 43 |
| S3$_A$ or I3$_A$ | A*A*A*T*T X A*A*A*A*A*A*A | 2 ± 2 | 19 ± 17 |
| S4$_A$ or I4$_A$ | A A A T T*X*A A A A A A A | 85 ± 16 | 1,830 ± 490 |
| S5$_A$ or I5$_A$ | A*A*A*T T X A A*A*A*A*A*A | 9 ± 4 | 55 ± 43 |
| S6$_A$ or I6$_A$ | A A A T*T*X*A*A A A A A A | 39 ± 9 | 400 ± 140 |

* = Phosphorothioate, dZ = 2′-deoxy-zebularine, All sugars are 2′-deoxyribose

**Table S2: Microscale Thermophoresis binding affinity values including uncertainties for A3A-targeting linear PS modified substrates and inhibitors**

| | Sequence (5′-3′) | Binding affinity (nM) |
|---|---|---|
| Substrate[a] | T T C A A | n/a |
| Inhibitor[a]: | | |
| H1$_A$ | T G C G C T T dZ G C G C A | 5.2 ± 1.8 |
| H2$_A$ | T*G*C*G*C*T*T*dZ*G*C*G*C*A | 9.1 ± 4.2 |
| H3$_A$ | T*G*C*G*C*T*T dZ G*C*G*C*A | 0.5 ± 0.1 |
| H4$_A$ | T*G*C*G*C*T*T dZ G*C*G*C*A | 1.1 ± 0.3 |

* = Phosphorothioate, dZ = 2′-deoxy-zebularine, ● = DNA, ● = LNA

**Table S3: Microscale Thermophoresis binding affinity values including uncertainties for A3A-targeting hairpin PS modified inhibitors**

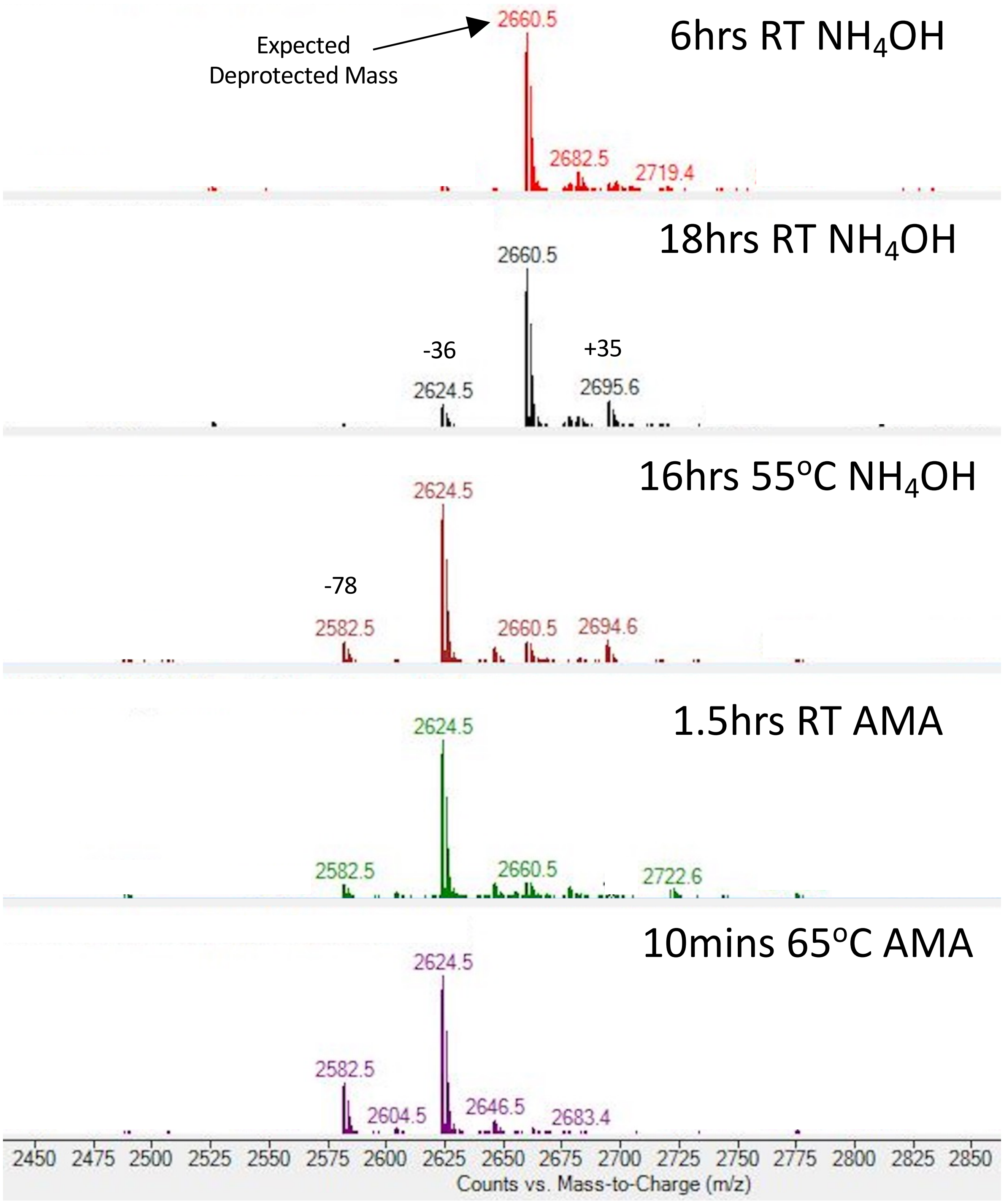

**Figure S1: Varying oligonucleotide deprotection conditions of I1$_G$ inhibitor (AATCCdZAAA) showing base-sensitivity of the dZ monomer. (AMA = 1:1 ammonium hydroxide:methylamine)**

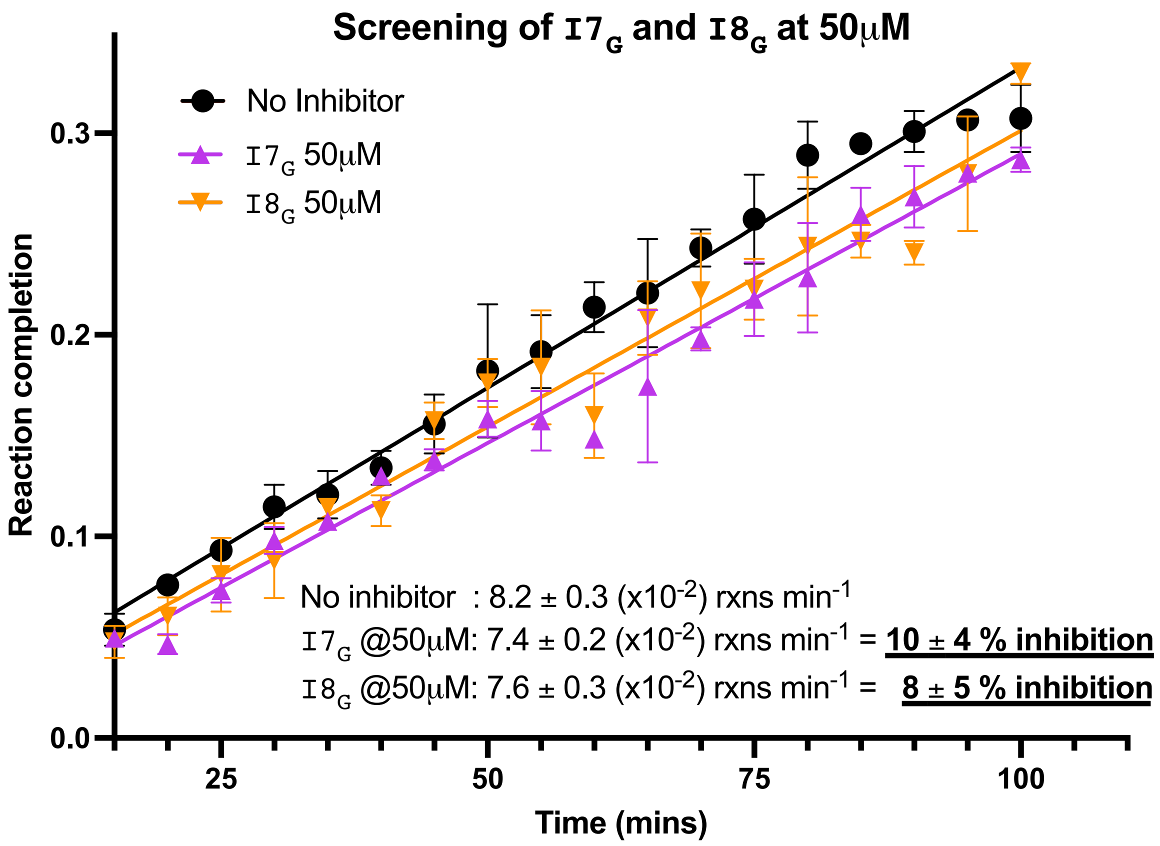

**Figure S2: Inhibition screening of S6 and S7 at 50μM against A3G-CTD2**

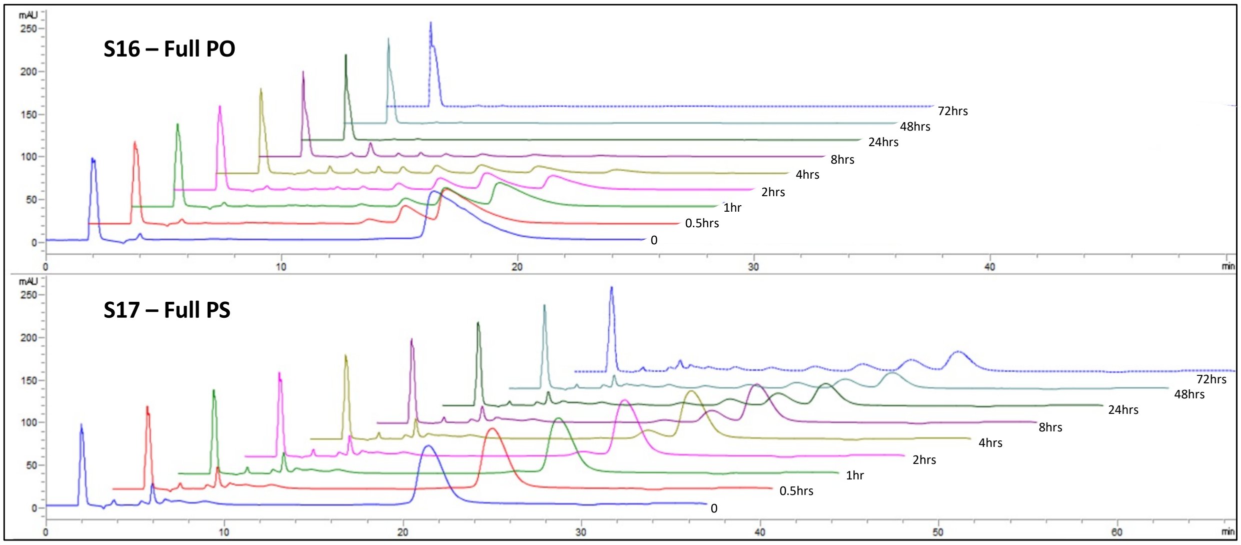

**Figure S3: Representative HPLC traces from nuclease stability assay. Linear A3A inhibitor S16 (full PO, top) compared to S17 (full PS, bottom) over multiple timepoints. Relative peak areas of the remaining full-length product were normalized and quantified to form data in Main Figure 6.**

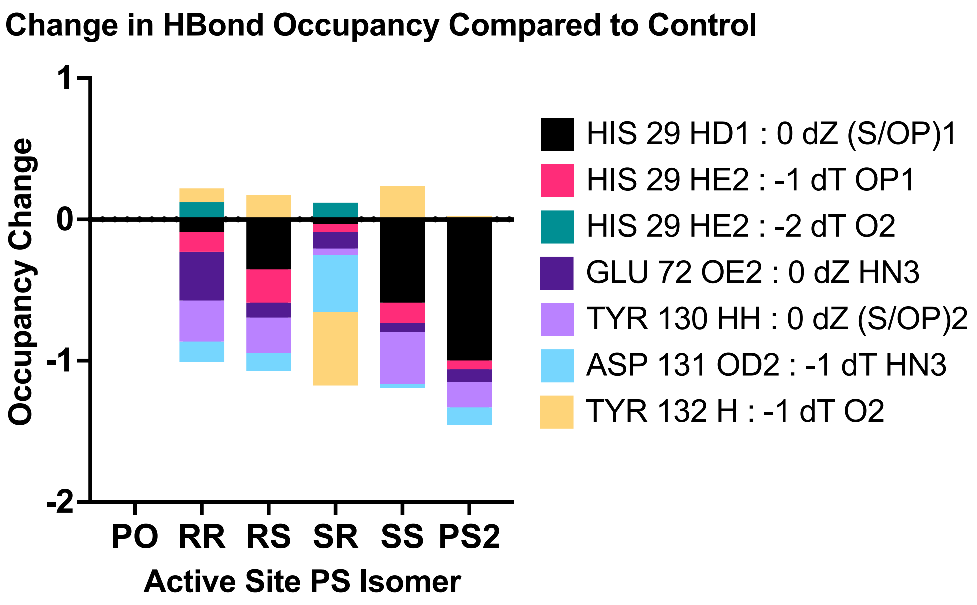

**Figure S4: Hydrogen bond occupancy changes relative to I1$_A$ for phosphorothioate (PS) or phosphorodithioate (PS2) flanking the target dZ in A3A targeting inhibitor (AAATTdZAAAAAAA). Data from 200 ns triplicate pMD simulations**

We considered the importance of chirality of PS in the positions flanking the active site, but computational modelling of the 4 diastereomeric PS combinations (i.e. *RR*, *RS*, *SR*, and *SS*) looked to be unfavored in all combinations compared to PO for A3A (**Fig S4**), likely due to the tight constraints of the active site. We also modelled achiral phosphorodithioate (PS2) linkages in these positions and found them also predicted to be disfavored, suggesting steric issues of the larger sulfur atoms around the tightly packed active site may be a problem. Thus we did not pursue these modifications experimentally.

**Protein Expression and Purification of A3G-CTD2**

*E. coli* BL21 cells transformed with a fusion pGEX construct of GST-A3G-CTD2 was cultured overnight at 37 °C with shaking in the presence of 100 μg/mL of ampicillin. The overnight culture was sub-cultured in fresh LB media at a 1:50 dilution and grown to mid log at 37 °C prior to induction with 1 mM IPTG. Protein expression occurred overnight at 18 °C. Cells were harvested by centrifugation, washed, and stored at -80 °C for protein purification. Overexpressed *E. coli* BL21 cells were resuspended in lysis buffer (50 mM sodium phosphate pH 7.3, 150 mM, 50 μM zinc chloride, 0.002 % Tween-20, and 1 mM DTT) and sonicated on ice. To obtain the soluble fraction, the cell lysate was clarified by two consecutive rounds of centrifugation at 30,000 x g for 30 minutes at 4 °C followed by syringe filtration using a 0.45 μm filter. The soluble fraction was then loaded onto a 5 mL GSTrap FF column (Cytiva), the column was washed with 50 mL of lysis buffer, and protein was eluted using GST elution buffer (50 mM sodium phosphate pH 7.3, 100 mM NaCl, 10 μM zinc chloride, 0.002 % Tween-20, 10 mM L-glutathione, and 1 mM DTT). GST cleavage was performed overnight by dialysis with PreScission Protease (Cytiva) in enzyme cut buffer (50 mM sodium phosphate pH 7.3, 100 mM NaCl, 10 μM zinc chloride, 0.002 % Tween-20, and 1 mM DTT). Isolation of native A3G-CTD2 was done using a reverse GST column run whereby native protein flows through the column while GST-A3G-CTD2 fusion protein and free GST is trapped. A3G-CTD2 was harvested in storage buffer (50 mM sodium phosphate pH 7.3, 150 mM NaCl, 50 μM zinc chloride, 0.002 % Tween-20, 10 % glycerol and 1 mM DTT) and frozen. The protein was quantified using the molar extinction 40,450 $M^{-1}$ $cm^{-1}$. Throughout the purification, sample purity was accessed by SDS-PAGE.
